# Resolving hematopoietic stem versus progenitor cell potential in the mouse dorsal aorta by differential *Runx1* +110 enhancer activity

**DOI:** 10.1101/2025.09.02.673630

**Authors:** Giorgio Anselmi, Vincent Frontera, Christina Rode, Andrew Jarratt, Naeema T. Mehmood, Matthew Nicholls, Stella Antoniou, Emanuele Azzoni, John Stamatoyannopoulos, Ditsa Levanon, Yoram Groner, Marella F.T.R. de Bruijn

**Affiliations:** MRC Molecular Haematology Unit, MRC Weatherall Institute of Molecular Medicine, Radcliffe Department of Medicine, University of Oxford, Oxford OX3 9DS, UK; Altius Institute for Biomedical Sciences, 2211 Elliott Ave, Seattle, WA 98121, USA; Department of Molecular Genetics, Weizmann Institute of Science, Rehovot, Israel; QIAGEN, Aix-en-Provence, France; Hulk Bio, West Linton, UK; Biobanking and BioMolecular Resources Infrastructure-European Research Infrastructure Consortium (BBMRI-ERIC), Graz, Austria; School of Medicine and Surgery, University of Milano-Bicocca, Monza Italy, and Fondazione IRCCS San Gerardo dei Tintori, Monza, Italy

**Author notes:** These authors contributed equally to this work (Giorgio Anselmi, Vincent Frontera, Christina Rode, Andrew Jarratt).

**Keywords:** hematopoietic stem cells, endothelial to hematopoietic transition, fate decisions, embryo, Runx1, enhancers

## Abstract

Hematopoietic stem cells (HSCs) are important in cell-based therapies for blood-related disorders. While progress has been made in the directed differentiation of pluripotent PSCs, such cultures promote hematopoietic progenitor cells (HPCs) over HSCs. Thus, elucidating signals, factors, and markers associated with HSC versus HPC lineage development in the embryo is imperative. During mouse embryonic development, HSCs and HPCs originate from hemogenic endothelium (HE) through a process critically dependent on the transcription factor Runx1. Here, we identified a *Runx1* enhancer that distinguishes emerging dorsal aorta HSCs from HPCs. Phenotypic, functional, and transcriptomic analyses of *Runx1* +110 enhancer-GFP reporter (110GFP) transgenic embryos showed that 110GFP expression marks HPCs, but not the emerging HSC lineage. Comparative transcriptomics revealed a 17-gene signature associated with *in vivo* long-term HSC potential. Furthermore, 110GFP- preHSCs showed increased expression of *Jarid2* and other PRC2 components, suggesting a role for epigenetic regulation in establishing the HSC fate during EHT. Finally, single-cell multiome analysis of dorsal aorta EHT identified the *Runx1* +3 enhancer as preferentially accessible in preHSC and underlined the specific activity of the +110 enhancer in HPCs. Our study demonstrates the power of cell-type specific enhancer-reporter models to dissect cell fate decisions in development and provides new inroads to label and/or perturb HSC versus HPC fate decisions *in vivo* and *in vitro*.

## INTRODUCTION

Hematopoietic stem cells (HSCs) are responsible for lifelong maintenance of hematopoiesis. They are clinically important in cell-based therapies for blood disorders. In spite of recent breakthroughs using chemically-defined cultures, it remains challenging to generate HSCs *de novo* in the laboratory^1^. To date, pluripotent stem cell-derived hematopoietic differentiation cultures generate many hematopoietic progenitor cells (HPCs) but only few HSCs^2,3^. During mouse embryonic development the hematopoietic system is established in multiple asynchronous waves, tightly restricted in space and time (reviewed in ref.^4^). The first two waves occur in the yolk sac (YS) between E7.25 and E9.5 and generate primitive erythrocytes, megakaryocytes, macrophages followed by definitive-type lineage-restricted HPCs^5^ ^6–8^. The third and last wave develops intra-embryonically in the midgestation aorta-gonad-mesonephros region (AGM) and is characterised by the generation of HSCs^9,10^. There, the HSC lineage emerges in the ventral wall of the dorsal aorta^11,12^ via a stepwise process in which they segregate from HE starting at E9.5 with the generation of proHSCs and mature through E10.5-E11.5 preHSC type 1 and type 2 intermediates into fully functional HSCs^13^. This endothelial-to-hematopoietic transition (EHT)^14^ is accompanied by the appearance of intra-aortic hematopoietic cell clusters that bud from the ventral wall of the dorsal aorta^14^. These clusters are heterogenous and contain (pre)HSCs as well as HPCs^15,16^. Between E10.5 and E11.5 the mouse aorta contains approximately 500-700 cluster cells. Only 1-2 of these are *bona fide* HSCs, as defined by their ability to sustain long-term serial multilineage engraftment in lethally irradiated mice^9,17,18^. Arguably, a better understanding of the regulatory events driving the establishment of HSCs versus HPCs from dorsal aorta HE, and when these potentials segregate, will aid the *de novo* generation of HSCs for therapeutic purposes.

To date, there is a paucity of reliable phenotypic markers discriminating between the emergence of HSC versus HPC potential during AGM EHT. To obtain such tools, we probed the *Runx1* locus for cell type specific cis-regulatory elements that could discriminate between cells with HSC and HPC potential as they emerge from HE. The transcription factor Runx1 is a master regulator of developmental hematopoiesis, with loss of Runx1 leading to a block in EHT at the level of the HE and a complete absence of all HSCs and HPCs (apart from primitive erythrocytes)^19–25^. The *Runx1* gene spans ∼220kb on mouse chromosome 16 and is transcribed from two alternative promoters, the distal P1 and proximal P2, which in isolation do not convey tissue-specific expression, suggesting that tissue/cell type specificity is mediated by distal enhancers^26–28^. Analysis of dynamic chromatin conformation changes during *in vitro* hematopoietic development suggested that the enhancers involved in mediating hematopoietic-specific *Runx1* expression are restricted to the gene body^29^. We previously identified and characterized the *Runx1* +23 enhancer, located in the first intron, which in transgenic enhancer-reporter mouse models mediates reporter gene expression to sites of hematopoietic cell emergence and colonization, in a spatiotemporal pattern that recapitulates endogenous Runx1 expression^30–33^. However, the +23 enhancer mediated reporter expression to all cells undergoing EHT, from (pre)HE to functional HSCs and HPCs^30,33^. To date, no *Runx1* cis-elements specific to either HE, HSCs, or HPCs have been reported.

Here, we report the specificity of the *Runx1* +110 enhancer in developmental hematopoiesis. We generated a transgenic mouse line carrying a GFP reporter transgene under the spatiotemporal control of the *Runx1* +110 enhancer. Phenotypic, functional, and transcriptomic analyses of GFP-expressing cells showed that the +110 enhancer recapitulates part of the endogenous *Runx1* expression pattern in yolk sac and AGM, labelling HPCs. Strikingly, however, no HE nor functional HSCs were marked by 110GFP expression. Leveraging the cell type specificity of 110GFP we identified a 17-gene signature that is specifically associated with long-term reconstituting HSC potential and present data suggesting an epigenetic regulation of establishing this potential. Finally, we generated a single cell Multiome dataset enriched for cells undergoing EHT, which corroborates the specificity of the *Runx1* +110 enhancer and identified an additional, complementary *Runx1* enhancer that is accessible in preHSC.

## RESULTS

### The *Runx1* +110 enhancer mediates reporter gene expression to distinct subsets of *Runx1*-expressing hematopoietic cells at the onset of hematopoiesis

Within the *Runx1* gene body, we have identified four enhancer regions which mediated *LacZ* reporter gene expression to the midgestation dorsal aorta where *Runx1* is expressed in cells undergoing EHT^30,34^ (Figure 1a). Analysis of transient transgenic concepti at E8.5 suggested that of these, the *Runx1* +110 enhancer displayed specificity for just part of the *Runx1* expression pattern, labelling hematopoietic cells in the yolk sac (YS) blood islands, but no endothelium in the paired dorsal aorta (Figure 1b). To comprehensively characterise the hematopoietic specificity of the *Runx1* +110 enhancer during the onset of developmental hematopoiesis, we generated a mouse line carrying a *hsp68Gfp+110* transgene, referred to as 110GFP (Figure 1c). Confocal microscopy and flow cytometry analysis of E8 110GFP transgenic concepti showed GFP expression in the YS blood islands, with 24.4±5.1% of Ter119+ erythrocytes (Ery) and 62.7±7.6% of VECad+Ter119-CD41+CD45-/+ hematopoietic progenitor cells (HPCs) expressing 110GFP, while virtually no 110GFP expression was seen in VECad+Ter119-CD41-CD45-endothelial cells (ECs) (Figure 1d,e and Figure S1a). This was also borne out functionally, with most YS CFU-Cs marked by the 110GFP transgene (Figure S1b).

**Figure 1.**
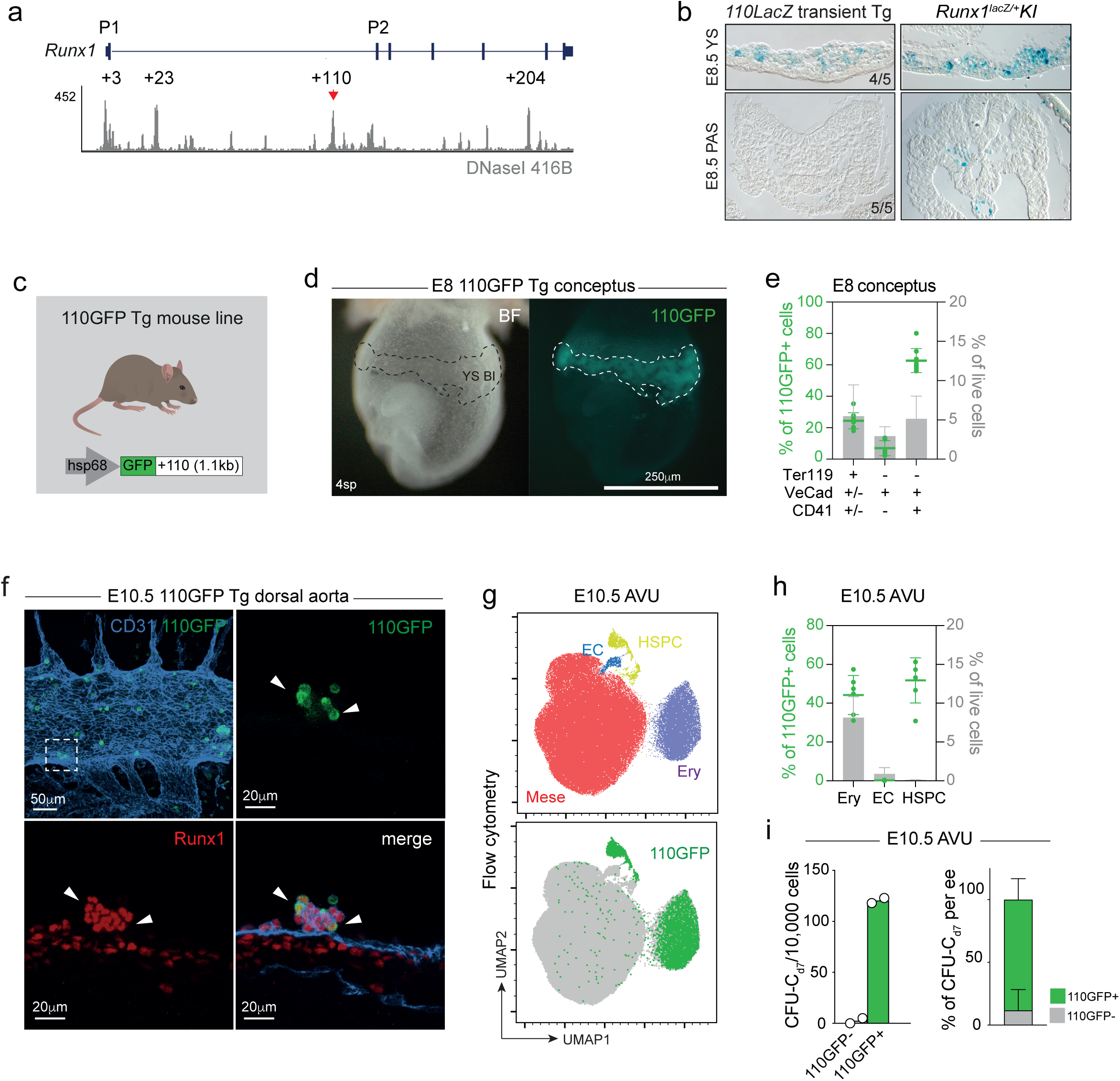
The *Runx1* +110 mediates reporter gene expression to distinct subsets of *Runx1*-expressing hematopoietic cells at the onset of hematopoiesis. **a.** DNaseI-seq track of the Runx1 locus in 416B cells^86^. The +110 *Runx1* enhancer is indicated with red arrowheads. **b.** Representative images of x-gal-staining (blue) in sections of E8.5 yolk sac (YS) and para-aortic splanchnopleura (PAS) from *Runx1* +110 enhancer-*lacZ* transient transgenic embryos. Runx1^lacZ/+^ KI embryos served as comparison. Fraction of embryos showing lacZ expression in specific tissues is reported in each image. **c.** Schematic of 110GFP enhancer-reporter generation. **d.** Whole mount image of 110GFP transgenic embryo at E8 (4sp). Dashed line highlights 110GFP expression in yolk sac (YS) blood island (BI). **e.** Quantification of 110GFP expression in E8 concepti (5-7sp) by flow cytometry (gating strategy in Figure S1b). Frequency ± SD of n=9 embryos in 2 independent experiments. **f.** Confocal whole mount immunofluorescence of E10.5 (36 sp) 110GFP transgenic embryo stained for CD31 (blue), Runx1 (red) and 110GFP (green). Solid arrowheads indicate examples of CD31+Runx+ hematopoietic clusters. **g.** UMAP plots of flow cytometry data from E10 AVU of 110GFP transgenic embryos. Cell type annotations based on marker expression (Figure S1d). GFP+ cells (green) are overlayed on GFP-cells (grey). Analysis performed on pooled tissues from n=3 110GFP embryos (33-34sp). **h.** Quantification of 110GFP expression in E10.5 (28-35sp) AVU by flow cytometry (gating strategy in Figure S1e). Frequency ± SD of pooled tissues from 6 independent experiments. **i.** CFU-C potential of GFP+ and GFP-cells isolated from AVU of E10.5 (32-34sp) 110GFP transgenic embryos. Absolute number of CFU-C_d7_ per 10,000 cells (n=2) and frequency of CFU-C_d7_ per embryo equivalent (ee) (mean of n=2 110GFP samples, 76.5% to 100% GFP+ range) are reported.

Analysis of the definitive hematopoietic wave in the midgestation dorsal aorta showed that, strikingly, 110GFP expression was restricted to part of the Runx1+ intra-aortic hematopoietic cell clusters budding into the vessel lumen (Figure 1f). This observation was corroborated by flow cytometry analysis of the E10.5 110GFP AGM+vitelline and umbilical arteries (AVU), which indicated that 110GFP-expressing cells mapped exclusively to the hematopoietic cell clusters, namely the immunophenotypically defined erythroid and hematopoietic stem/progenitor cell (HSPC) clusters (Uniform Manifold Approximation and Projection (UMAP) in Figure 1g and Figure S1c). Further quantification showed that 110GFP labelled 44.2±10.1% of erythrocytes and 51.7±11.6% of CD41+CD45+ HSPCs in the E10.5 AVU (Figure 1h and Figure S1d). In agreement with the immunophenotypic data, functional analysis of 110GFP+ cells isolated from the midgestation AVU showed that over 70% of CFU-Cs were marked by the transgene (Figure 1h). Altogether, this demonstrates that *Runx1* +110 enhancer mediated reporter gene expression is restricted to part of the hematopoietic cells at intra- and extra-embryonic sites of HSPC emergence.

### The *Runx1* 110GFP enhancer-reporter marks primitive macrophages, EryP and emerging *Hoxa+* HSPCs

To comprehensively identify the hematopoietic cells labelled by the 110GFP transgene, 110GFP+ cells were isolated from the AVU of E10 (31-34sp) embryos and processed for scRNAseq (10x Genomics; Figure 2a). In parallel, a reference dataset that captured all AVU hemogenic and hematopoietic cells was generated by profiling GFP+ cells isolated from E10 (32-35sp) 23GFP transgenic embryos, as we previously showed that the 23GFP enhancer-reporter labels all cells undergoing EHT, from the HE through to the emerging HPCs and HSCs^32,33^. After quality control, the 110GFP+ and 23GFP+ datasets were integrated and non-hematopoietic contaminants removed (Figure S2a-c). In the UMAP of the remaining 2,260 110GFP+ and 4,246 23GFP+ cells twelve distinct cell clusters were identified, covering EHT and differentiating hematopoietic lineages (Figure 2b; Figure S2d, e; Table S3). As expected, the 23GFP reporter labelled cells in 11 out of the 12 UMAP clusters, i.e. all but the EryP as 23GFP captures all Runx1+ hematopoietic cells (EryP are Runx1/23GFP-negative at this timepoint)^22,30^. In contrast, 110GFP labelled predominantly the macrophage lineage clusters pMac, Mac1 and Mac2, as well as EryP but virtually none of the definitive erythrocytes (EryD cluster; Figure 2b, c) and other lineage progenitors (MEP, Mk, pMC and pNeu, Figure S2f, g). In addition, 110GFP labelled HPCs (cluster HPC1, HPC2), but very few if any ECs (cluster Endo; Figure 2b), in line with the immunophenotypic and functional analyses (Figure 1f, g, h). Finally, 110GFP labelled some cells in the UMAP cluster with EHT characteristics, i.e. expressing endothelial genes together with *Runx1*, *Hlf* and Myb (Figure 2b, d). Of note, only the Endo, EHT, and HPC1 clusters expressed *Hoxa* genes (Figure 2d, e and Figure S2h), indicative of a recent intraembryonic origin^35,36^.

**Figure 2.**
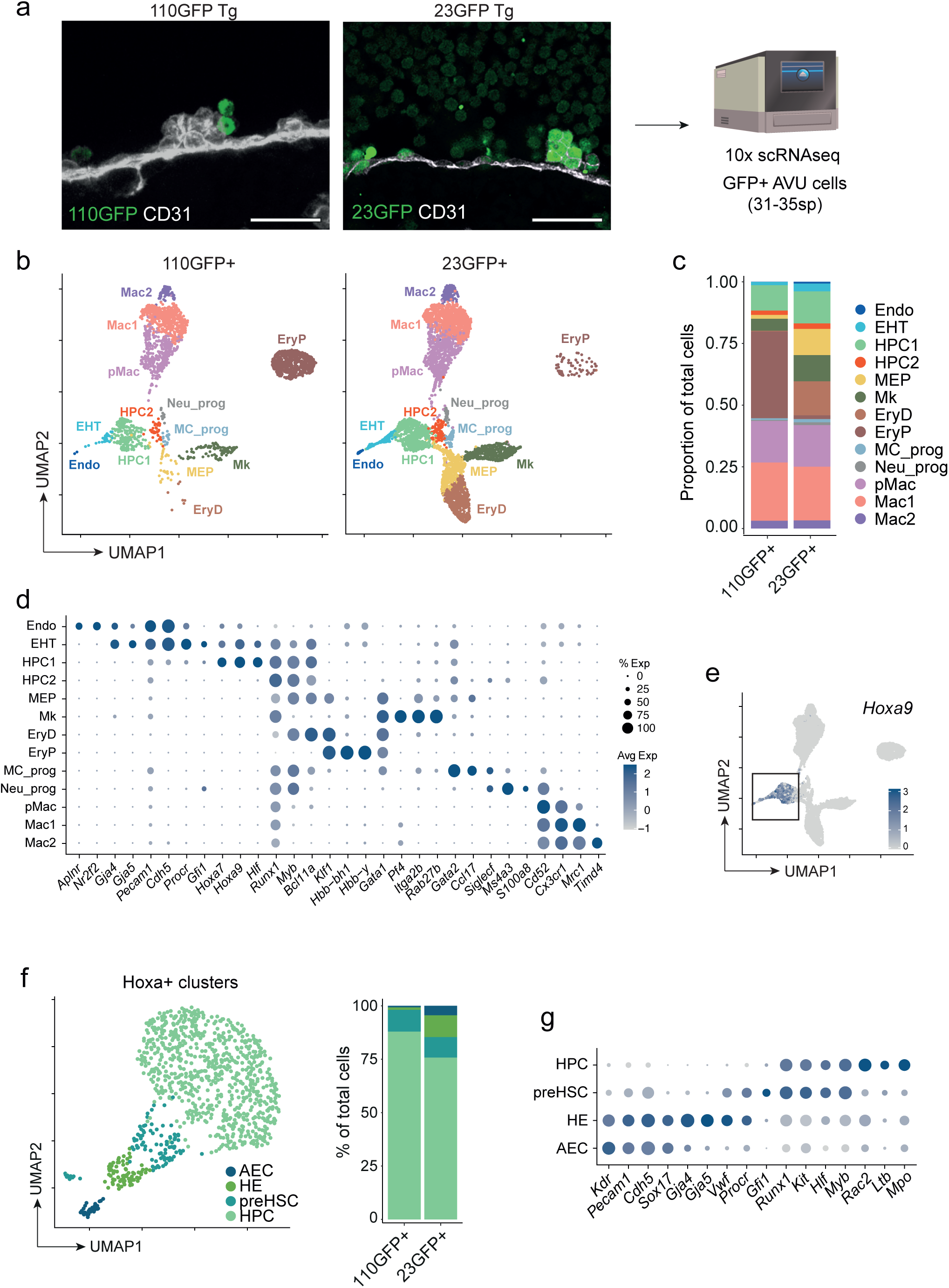
The 110GFP transgene marks primitive macrophages, EryP and emerging *Hoxa+* HSPCs. **a.** Schematic of scRNA-seq analysis (Chromium 10x) of E10 110GFP+ (31-34sp) and 23GFP+ (32-35sp) cells from AVU (pooled tissues from n=7 and n=9 embryos, respectively). **b.** UMAP representation of transcriptomic profiling of 110GFP (2,997) and 23GFP (5,751) cells from E10 AVU. Unsupervised clustering resulted in 13 clusters named according to the expression of cell type-specific marker genes (Figure 3d). Endo, endothelial cells; EHT, endothelial-to-hematopoietic transition; HPC, hematopoietic progenitor cells; MEP, megakaryocytes and erythrocytes progenitor; Mk, megakaryocytes; EryD, definitive erythrocytes; EryP, primitive erythrocytes; MC_prog, mast cell progenitor; Neu_prog, neutrophil progenitor; pMac, macrophage precursors; Mac, macrophage. **c.** Bar graph displaying the frequency of clusters within each sample. **d.** Dot plot displaying marker genes defining cell clusters in Figure 3b. **e.** Expression of *Hoxa9* in 110GFP+ and 23GFP+ integrated dataset (Figure 2b). **f.** UMAP representation and unsupervised clustering of a subset of cells expressing *Hoxa7*, *Hoxa9* and *Hoxa10* genes (top left), with bar graph displaying the frequency of clusters within each sample. **g.** Dot plot of marker genes used to annotate *Hoxa+* clusters. AEC, arterial endothelial cells; HE, hemogenic endothelium; preHSC, HSC precursors; HPC, hematopoietic progenitor cell (bottom).

To further examine the cells labelled by 110GFP among the emerging *Hoxa+* intraembryonic cell populations, we reclustered the Endo, EHT, and HPC1 UMAP clusters, which resulted in the identification of 4 main subclusters (Figure 2f). Based on differential endothelial and hematopoietic gene expression these were annotated as an arterial endothelial cluster (AEC) expressing *Cdh5*, *Sox17* and *Gja4*, a HE cluster expressing arterial genes (*Gja4* and *Gja5*), low levels of *Runx1*, and other hematopoiesis-associated genes (*Vwf* and *Procr*), a HPC cluster expressing canonical hematopoietic markers such as *Runx1*, *Kit*, *Rac2* and *Mpo,* and a cell cluster characterised by the expression of *Vwf*, *Procr*, *Gfi1* along with *Runx1*, *Kit* and *Hlf*, and absence of arterial gene expression, annotated as preHSC (Figure 2f,g Table S4). Comparison with published endothelial and hematopoietic gene module scores^37–39^ further supported this annotation (Figure S2i). While the AEC and HE UMAP clusters were uniquely present among 23GFP+ cells, the preHSC and HPC clusters were present among both 110GFP+ and 23GFP+ AVU cells (Figure 2f, bar graph), raising the question whether the HSC lineage is labelled by the *Runx1* +110 enhancer-reporter.

### Cells belonging to the functional HSC lineage are not marked by the 110GFP reporter

Flow cytometric analysis of the cells of the HSC lineage showed that virtually none (4.5±1.1%) of the E9.5 proHSCs (VEC+Kit+CD41+CD43-CD45-) in the PAS+VU expressed 110GFP (Figure 3a, b). In contrast, 38.8±4.1% of phenotypic E10.5 preHSC1 (VEC+Kit+CD41+CD43+CD45-) and 50.5±6.5% of phenotypic E11.5 preHSC2/HSCs (VEC+CD45+CD43+Sca1+) were marked by 110GFP reporter expression (Figure 3a, b). Of note, while HPCs can be separated from E9.5 pro-HSCs on the basis of CD43 expression (with just under half (45.8±3.9%) of the phenotypic E9.5 HPCs (VEC+Kit+CD41+CD43+CD45-) expressing 110GFP; Figure S3a), both E10.5/11.5 preHSCs/HSCs and HPCs of the AVU express CD43^13^, thus requiring functional tests to assess whether preHSCs and HSCs express 110GFP. This was done by transplantation into irradiated adult recipients either directly (for HSCs) or after OP9 co-aggregate culture (for preHSCs^13,40^). Our results clearly showed that functional preHSCs and HSCs potential segregated entirely to the 110GFP-AVU cell populations (Figure 3c, d). In line with this, phenotypic-defined LT- and ST-HSCs in adult BM also lacked 110GFP expression (Figure S3b). Interestingly, a subset of phenotypic HSCs (CD48-CD150+/- LSK) in the E14 foetal liver did express 110GFP (Figure S3c), which may be due to this population also containing hematopoietic cells devoid of LT-HSC potential^41^.

**Figure 3.**
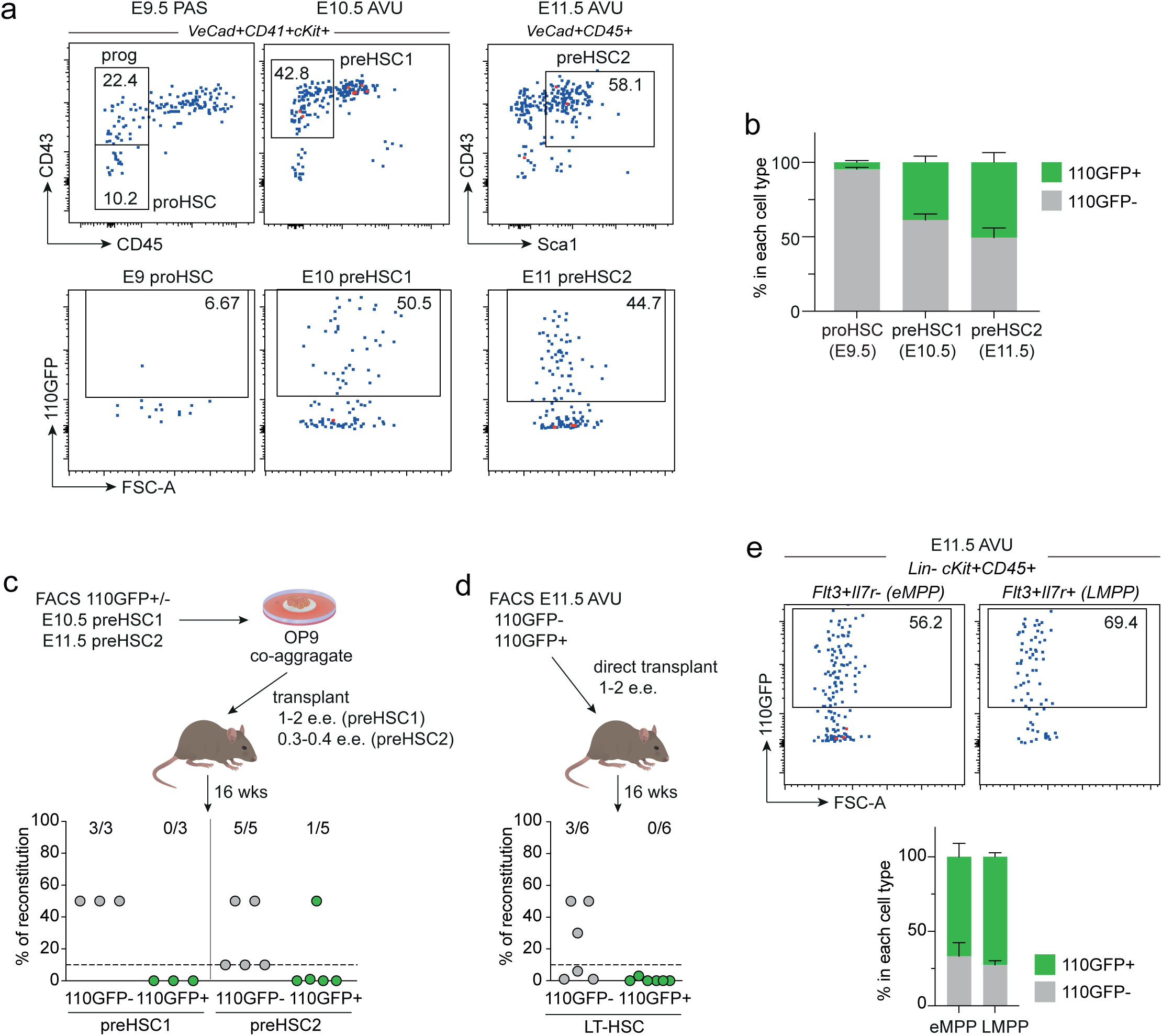
Cells belonging to the functional HSC lineage are not marked by the 110GFP reporter. **a.** Flow cytometry analysis of 110GFP expression in E9.5 (18-25sp) pro-HSCs (pooled PAS from n=3 independent experiments), E10.5 (28-36sp) preHSCs type1 (pooled AVU from n=6 independent experiments) and E11.5 (12-18tsp) preHSCs type2 (pooled AVU from n=5 independent experiments). **b.** Quantification of cell types marked by 110GFP expression in Figure 3a. Mean ± SD of pooled tissues from n=3 E9.5, n=6 E10.5 and n=5 E11.5 independent experiments. **c.** Repopulation activity of 110GFP+ or 110GFP-pre-HSCs type1 (1-2ee transplanted in n=3 mice from 1 independent experiment) and pre-HSCs type2 (0.3-0.4ee transplanted in n=5 mice from 2 independent experiments) after OP9 co-aggregate cultures. Peripheral blood reconstitution was assessed by genomic PCR 16 weeks post-transplant. Engraftment threshold 10% (dotted line). Number of reconstituted mice out of the total transplanted animals is reported. **d.** Repopulation activity of 110GFP+ and 110GFP-cells isolated from E11.5 AVU (1-2ee transplanted). Peripheral blood was analysed by genomic PCR 16 weeks post-transplant. Reconstitution threshold 10% (dotted line). Data show the number of reconstituted/transplanted animals (n=6 from 4 independent experiments). **e.** Flow cytometry analysis of 110GFP expression in phenotypically defined eMPP (lin-CD45+cKit+Flt3+Il7r-) and LMPP (lin-CD45+cKit+Flt3+Il7r+) from E11.5 (15-20tsp) AVU. Frequency ± SD of n=8 independent experiments.

A recent study suggested that AGM-derived preHSCs may give rise to embryonic multipotent progenitors (eMPPs) that, independent of HSCs, are responsible for a significant fraction of multilineage adult blood^42^. These eMPPs were identified as Flt3-expressing cells within E10.5-E11.5 cKit+CD45+Il7r-hematopoietic cells and they were shown to contribute significantly to adult MMP3/4^42^. Analysis of 110GFP expression among Kit+CD45+Flt3+Il7r-cells in the E11.5 AVU showed that approximately two-thirds (66.7±9%) of these phenotypic eMPPs expressed 110GFP (Figure 3e, Figure S3d). While expression of 110GFP in part of the MMP3 and MMP4 in E14 foetal liver and adult bone marrow (Figure S3b, c) is suggestive of 110GFP labelling functional eMPPs, examining this directly will require lineage tracing experiments beyond the scope of this study. Finally, also part of the Kit+CD45+Flt3+Il7r+ lymphoid-primed multipotent progenitors (LMPP) expressed 110GFP (72.5+2.8%; Figure 3e, Figure S3d). In summary, our data showed that functional HSCs and preHSCs of the midgestation embryo are not marked by 110GFP, while the progeny of (pre)HSCs may be labelled.

### Emergence of HSC potential in 110GFP-cells is marked by a 17-gene signature and expression of genes associated with epigenetic repression

Next, we sought to identify markers that could distinguish cells with preHSC potential among all emerging hematopoietic cells in the midgestation AVU. Analysis of known HSC-associated genes in the 23GFP+ versus 110GFP+ *Hoxa+* ‘preHSC’ UMAP cluster (Figure 2f) showed some genes with distinct expression patterns in 23GFP+ versus 110GFP+ cells (Figure S4a). For example, *Tcf15, Pdzk1ip1* and *Id3,* previously associated with adult BM LT-HSC activity^43–45^, were expressed among 23GFP+ preHSCs (Figure S4a), in line with 23GFP marking the HSC lineage throughout development^32,33^ (Figure S4b). In contrast, these HSC-associated genes were low/absent from the 110GFP+ ‘preHSC’ cluster, in line with the observed lack of functional preHSCs (Figure S4a; Figure 3c). Similarly, *Cd27*, reported to be expressed at intermediate levels in HSCs^39^, was expressed at low/intermediate levels in 23GFP+, but at high levels in 110GFP+ cells (Figure S4a). Thus, based on differential expression of HSC-associated genes between 110GFP+ and 23GFP+ ‘preHSC’ UMAP cluster cells, we divided the ‘preHSC’ UMAP cluster into two distinct subsets, where one harbours functional preHSCs (preHSC^Pot^) and the other does not (preHSC^Not^) (Figure S4d and Figure 4a). While at the protein level preHSC^Pot^ and preHSC^Not^ cells were indistinguishable based on expression of cell membrane markers for HSPCs and lineage committed progenitors (CITEseq for CD41, CD45, cKit, CD127, CD115; Figure S4e), direct comparison and differential gene expression testing of preHSC^Pot^, preHSC^Not^, and HE cells (see Methods), identified a 17-gene signature that highly correlated with preHSC potential (Figure 4b,c, Table S5). Of note, this signature did not include some known HSC-associated genes, as they were also highly expressed in HE (e.g. *Tcf15* and *Mllt3*, Figure S4f). Importantly, this 17-gene signature was enriched also in functionally validated preHSC and HSC in publicly available datasets^37,39^ (Figure S4f-i), indicating its broader applicability in distinguishing HSC lineage potential.

**Figure 4.**
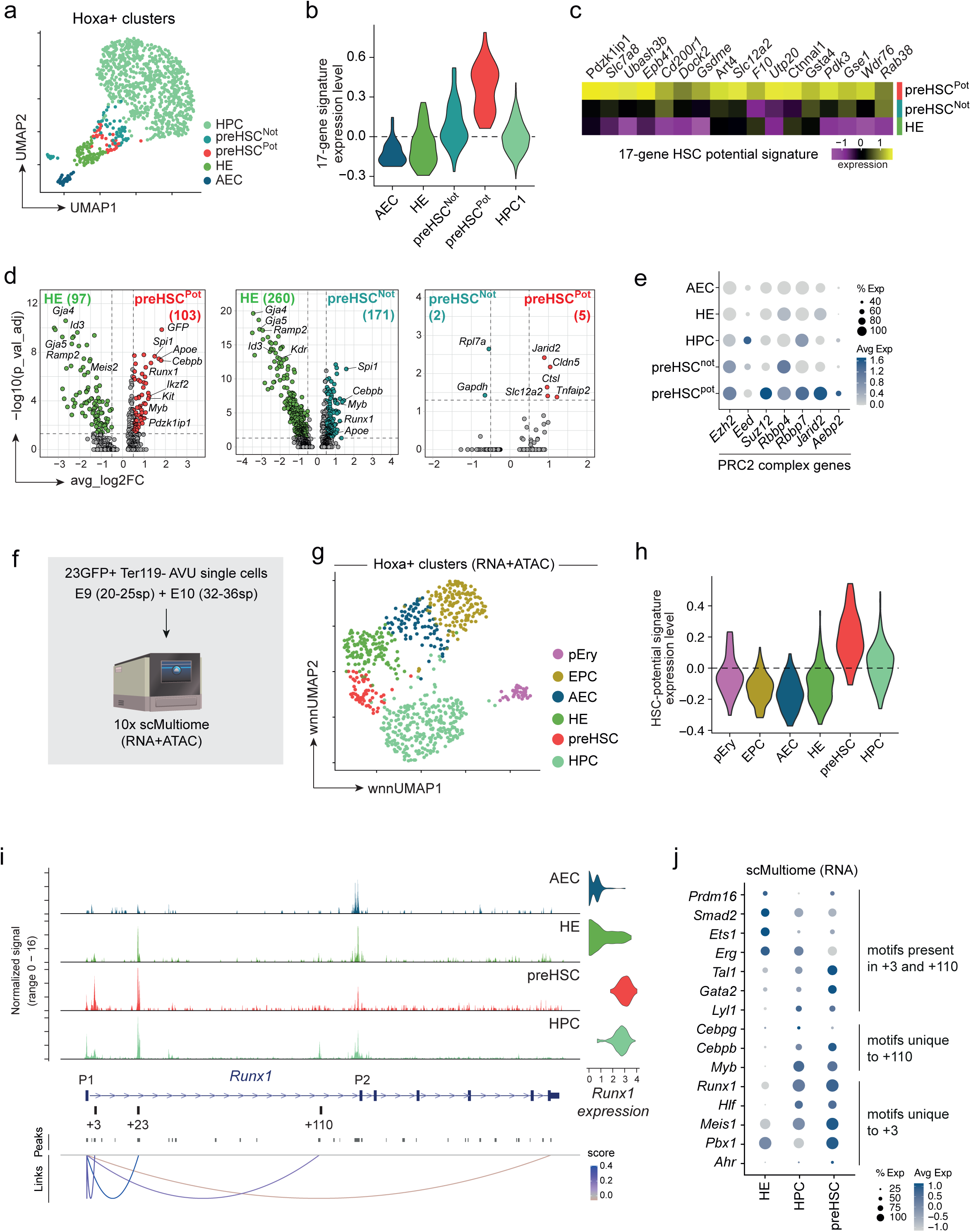
Emergence of HSC potential in 110GFP-cells is marked by a 17-gene signature and expression of genes associated with epigenetic repression. a. UMAP visualization of *Hoxa+* clusters from Figure 2f, with refined annotation to include functionally defined preHSCs (as defined in Figure S4d). b. Expression of the HSC potential signature (17 genes) in refined-annotated Hoxa+ clusters, with the functionally defined preHSC^Pot^ cluster coloured in red. c. Heatmap displaying the HSC potential signature as calculated by 3-way comparison of HE, preHSC^Not^ and preHSC^Pot^ clusters. Each row represents a pseudobulk average of all cells from a single cluster. d. Volcano plot displaying differentially expressed genes (Wilcoxon Rank Sum test p<0.05, log2 fold change > 0.5) in pairwise comparisons of HE, preHSC^Pot^ and preHSC^Not^. e. Dot plot showing the average expression of the PRC2 complex genes in *Hoxa+* clusters defined in Figure 4a. f. Schematic of scMultiome analysis (Chromium 10x) of E9 (20-25sp) and E10 (32-36sp) Ter119-23GFP+ cells from AVU (pooled tissues from n=33 and n=24 embryos, respectively). g. UMAP representation of joint RNA and ATAC measurements using weighted nearest neighbour (WNN) integration for *Hoxa+* clusters from multiomic data. h. Expression of the HSC potential signature (17 genes) in *Hoxa+* clusters from multiomic data. i. ATAC-seq tracks and peak-gene links for the Runx1 locus in *Hoxa+* clusters. AEC, arterial endothelial cell; HE, hemogenic endothelium; preHSC, HSC precursor; HPC, hematopoietic progenitor cell. j. Dot plot displaying gene expression of representative transcription factors whose motifs were found in conserved phylogenetic footprints of the +3 and +110 enhancers.

To explore what molecular mechanisms could be underlying the establishment of HSC potential in the dorsal aorta, we performed pairwise comparisons between preHSC^Pot^, preHSC^Not^ and HE. Both preHSC^Pot^ and preHSC^Not^ were clearly distinct from HE, with significantly upregulated hematopoiesis-associated genes (e.g. *Runx1*, *Spi1*, *Cebpb*, *Kit* and *Myb*) and downregulated HE-associated arterial markers (e.g. *Gja4*, *Gja5*, *Id3* and *Ramp2)* (Figure 4d, Table S6-9). In contrast, only few differentially expressed genes could be identified between preHSC^Pot^ cells and preHSC^Not^ (Table S10-11), suggestive of a high degree of transcriptional similarity. Interestingly, the most significantly upregulated gene in preHSC^Pot^ was *Jarid2* (Figure 4d). Jarid2 is part of the Polycomb repressive complex 2 (PRC2) and in line with this, an increased expression of PCR2 core (*Eed*, *Suz12*, *Ezh2* and *Rbbp4* or *Rbbp7)* and accessory (*Aebp2)* subunits was observed in preHSC^Pot^ cells (Figure 4e).

### Identification of a *Runx1* enhancer active during AVU EHT that is complementary to the +110 enhancer

Having established that *Runx1* +110 enhancer activity anti-correlates with HSC emergence, we asked whether there are also *Runx1* enhancers with activity that positively correlates with the HSC lineage, while not active in HPCs. For this, we generated a single cell Multiome dataset (RNA+ATAC) on 23GFP+ cells undergoing EHT in the E9.5 and E10.5 AVU (Figure 4f, Figure S5a-b and Table S12). Re-clustering the *Hoxa*-expressing endothelial and EHT cell clusters (Figure S5c) identified six distinct populations (Figure 4g, Figure S5d-e). Four of these corresponded to AEC, HE, preHSCs and HPC also identified in the E10 scRNAseq (compare Figure 4f and 2f). The remaining two were endothelial progenitor cells (EPCs) and a small cluster expressing genes associated with early erythroid differentiation, labelled as erythrocyte progenitors (pEry) (Figure S5f). Our 17-gene HSC potential signature (Figure 4c) was enriched in the preHSC multiome cluster indicating the presence of cells with *bona fide* HSC potential (Figure 4h). Analysis of ATAC tracks in the AEC, HE, preHSC, and HPC cell clusters confirmed that the *Runx1* +110 enhancer is accessible in AVU HPCs only (Figure 4i), while the +23 enhancer is active in HE, preHSC and HPCs. Nevertheless, levels of *Runx1* gene expression were comparable between the preHSC and HPC multiome clusters (Figure 4i). Interestingly, a clear peak of increased chromatin accessibility, unique to the preHSC cluster, was seen at the *Runx1* +3 enhancer (Figure 4i and Figure S5g). Comparing candidate regulatory modules in the +3 and +110 enhancers showed that of the 35 identified TF motifs (filtered based on mean conservation score and expression of upstream TFs in the preHSC/HPC clusters, see Methods), 11 were shared while 13 and 11 were unique to the +3 and +110 enhancers, respectively (Table S13-14). Shared motifs included broad hematopoietic TF motifs such as ETS, E-box, GATA (Figure S6 and S7). Motifs unique to the +110 enhancer included CEBP and MYB (Figure S7), in line with the labelling of myeloid cells by 110GFP. The +3 enhancer, in contrast, harboured RUNX, HLF, MEIS, PBX, and AHR motifs (Figure S6). Interestingly, expression levels of upstream TF for these motifs differed between HE, preHSC and HPC (Figure 4j). In summary, during EHT the +3 and +110 enhancers show complementary chromatin accessibility patterns that together with the specific patterns of TF motifs, are indicative of distinct cell type specificity of these enhancers, and suggestive of the +3 marking the emerging HSC lineage.

## DISCUSSION

During embryonic development, the first HSCs are generated alongside HPCs in the major arteries of the embryo, the dorsal aorta and vitelline and umbilical arteries. Studies into the generation of these cells have proven instrumental in developing *in vitro* cultures aimed at generating HSPCs for research and therapeutic purposes. However, such cultures generate a mix of HSCs and HPC and are not yet selective for HSC production. Optimisation has proven painstakingly hard, in part due to a paucity of markers to specifically monitor HSC generation. Here, we show that *Runx1* +110 enhancer activity anti-correlates with the emergence of HSC potential in the major arteries of the mouse embryo and demonstrate its use in elucidating possible mechanisms involved in HSC specification *in vivo*.

In developmental hematopoiesis, the +110 enhancer-GFP reporter labelled only subsets of clonogenic HPCs in the yolk sac and dorsal aorta. In line with this, single cell transcriptome analysis of 110GFP+ AVU cells showed labelling in a population of HPCs, along with labelling of primitive erythrocytes and macrophages. In addition, 110GFP labelled cells phenotypically and transcriptionally defined as preHSC, but, strikingly, these contained no functional preHSCs or HSCs (preHSC^Not^). We exploited the absence of 110GFP labelling on dorsal aorta preHSC^Pot^ in a comparison of the 110GFP+ transcriptomic data with a highly enriched 23GFP+ EHT dataset we generated in parallel, to identify a 17-gene signature associated with the emergence of long-term HSC potential in the mouse embryo. This signature was also enriched in functionally validated dorsal aorta preHSC, HSC, as well as fetal liver and bone marrow HSCs in other scRNAseq datasets of mouse HSC development^37,39^. Of note, markers known to be associated with emerging HSCs, such as *Hlf*, *Mllt3*, *Mecom*^46^ were not part of the signature, as we found them to be expressed at comparable levels to preHSC in HE and/or HPCs. Instead, our signature included *Pdzk1ip1* (aka Map17) and *Ctnnal1* (aka α-catulin) that have been associated with adult BM HSCs^44,47–51^. Combination of an α-catulin-GFP reporter and c-kit expression has been used to specifically identify HSCs in mouse bone marrow^50,51^, further supporting the specificity of our 17-gene signature to identify HSCs/HSC-fated cells in the embryo. Thus, the *Runx1* +110 enhancer-reporter and the 17-gene HSC potential signature are valuable negative and positive indicators of HSC potential, respectively. It will be of interest to explore their use in designing and monitoring robust *in vitro* culture conditions aimed at de novo generation and/or expansion of HSCs.

We observed a higher expression of *Jarid2* in preHSC^Pot^ than in preHSC^Not^. Jarid2 is a member of the Polycomb repressive complex 2 (PRC2), a chromatin-biding complex involved in repressing gene expression through its histone methyltransferase activity^52^. Other components of the PRC2 complex, including the core subunits *Ezh2, Suz12, Rbbp7* and the accessory protein *Aebp2* were also expressed in preHSC^Pot^, suggesting a PRC2-mediated role of Jarid2 in nascent HSC development. The role of PRC2 during embryonic hematopoiesis, and more specifically in the specification of the HSC lineage, is not fully elucidated. Previous work has provided contradictory data on the impact of PRC2 inactivation, depending on the targeted PRC2 component and the Cre driver used in conditional knockout mouse models. Indeed, while an essential role of PCR2 in foetal HSC formation and function has been proposed^53–55^, other reports have suggested that Ezh2 is dispensable for the maintenance of FL HSC^56–59^. Jarid2 itself has been previously associated with regulation of differentiation and function in multiple stem cell compartments^60–62^ and it has been shown to act either by driving PRC2 activity on chromatin or in a PRC2-independent fashion^63^. The precise role of Jarid2 in preHSC^Pot^ requires further experimentation.

The transcriptional regulation of *Runx1* is complex and expression levels require precise spatiotemporal control for hematopoietic development to proceed normally and to maintain adult hematopoietic homeostasis^24,64,65^. To assess what enhancers apart from the +110 and +23 may be active during EHT, we analysed chromatin accessibility in primary E9.5-E10.5 dorsal aorta EHT cells (by scMultiome), isolated on the basis of 23GFP expression. These analyses corroborated the specificity of the +110 enhancer for HPCs and that of the +23 enhancer for all cells undergoing EHT. Interestingly, accessibility of another *Runx1* hematopoietic enhancer element, the +3, peaked in preHSCs, making it complementary to the +110 in EHT. TF motif analysis showed part overlapping and part specific motifs. A recent study by Frömel et al. on the design principles of cell-state-specific enhancers in hematopoiesis showed a complex interplay between cell state, site combinations, and TF occupancy determining the repressive or activating function of identical TF motifs^66^. Gata1/2, Cebpa, Meis1, Runx1 TFs were reported to be involved in such complex interplays that could lead to cell type specific activation, neutralization, or repression depending on the cell lineage. Interestingly, the +110 enhancer harbours both GATA and CEBP motifs, while the +3 enhancer harbours GATA, MEIS and RUNX motifs, with corresponding upstream TFs showing dynamic expression between HE, HPC and preHSC. This suggests that the specificity of these enhancers is likely determined by a complex interplay between these factors on the *Runx1* enhancer elements. Unravelling the full regulatory complexity of *Runx1* will be relevant to better understand *RUNX1*-associated hematopoietic disorders.

Altogether, our data support a working model in which a phenotypic and transcriptionally homogeneous pool of preHSC arises from HE in the major arteries of the embryo (in contrast to previous studies^67–69^, we did not observe transcriptional heterogeneity among HE). A subset of the preHSC, the preHSC^Not^ (in which both the *Runx1* +23 and +110 enhancer are active) will only contribute to HSC-independent hematopoiesis, including HPCs and possibly eMPPs. In contrast, the preHSC^Pot^ (in which the +23 and +3 enhancers are active), express high levels of *Jarid2* which we hypothesise locks them into a more quiescent state, allowing their maturation into engraftable HSC^70^. Future studies aimed at resolving the precise cellular origins and niches of preHSC^Pot^ and preHSC^Not^ will shed light on the molecular mechanisms determining these potentials and feed into designing and monitoring robust *in vitro* culture conditions aimed at *de novo* generation and/or expansion of HSCs.

## RESOURCE AVAILABILITY

### Lead contact

Requests for further information and resources should be directed to and will be fulfilled by the lead contact Marella F. T. R. de Bruijn.

### Material Availability

All unique/stable reagents generated in this study are available from the lead contact with a completed materials transfer agreement.

### Data and code availability

Single-cell RNA-seq and single-cell Multiome data are available under the ArrayExpress accession number xxxx and xxxx. This paper does not report original code. Any additional information required to reanalyse the data reported in this paper is available from the lead contact upon request

## ACKNOWLEDGEMENTS

We thank current and past members of the de Bruijn laboratory for their practical help, advice and discussions, Simone Riva for computational advice, and Pik-Shan Li and Jackie Sloane-Stanley for generation of mouse transgenics. This work was supported by programs in the MRC Molecular Haematology Unit Core award to MDB (MC_U137970202, MC_UU_12009/2, MC_UU_00016/2, MC_UU_00029/5) and a Weizmann - UK Joint Research Program (’Making Connections’) supported by the Weizmann friends in the UK to MDB, DL, YG. We would like to acknowledge Kevin Clark, Sally Clark, and Paul Sopp in the flow cytometry facility at the MRC WIMM for providing cell sorting services. The facility is supported by the MRC TIDU; MRC MHU (MC_UU_12009); NIHR Oxford BRC; Kay Kendall Leukaemia Fund (KKL1057), John Fell Fund (131/030 and 101/517), the EPA fund (CF182 and CF170) and by the MRC WIMM Strategic Alliance awards G0902418 and MC_UU_12025. We thank Neil Ashley and Maria Greco for help on 10x Chromium sample processing. The WIMM Single Cell Core Facility was supported by the MRC MHU (MC_UU_12009), The Oxford Single Cell Biology Consortium, (MR/M00919X/1) and the WT-ISSF (097813/Z/11/B#). The facility was supported by WIMM Strategic Alliance awards G0902418 and MC_UU_12025. We would like to acknowledge Tim Rostron in the Sequencing facility at the MRC WIMM for providing sequencing services. The facility is supported by the MRC TIDU and by the EPA fund (CF268). We thank Christopher Lagerholm and Jana Koth for their imaging assistance. The Wolfson Imaging Centre Oxford is supported by the MRC via the WIMM Strategic Alliance (G0902418), the MHU (MC_UU_12009), the TIDU (MC_UU_12010), the Wolfson Foundation (18272) and the Wellcome Trust (Micron 107457/Z/15Z) grants.

## AUTHOR CONTRIBUTIONS

Conceptualization: GA, MDB; Methodology: GA, VF, CR, AJ, NM, MN, SA, EA, MDB; Formal analysis: GA, VF, CR; Investigation: GA, VF, CR, DL, YG, MDB; Resources: GA, CR, AJ, JS, MDB; Data curation: GA; Writing – original draft: GA, CR, MDB; Writing – review & editing: GA, CR, MN, EA, DL, YG, MDB; Visualization: GA, CR, MN; Supervision: GA, MDB; Funding acquisition: DL, YG, MDB.

## SUPPLEMENTAL INFORMATION

Document S1. Figures S1-S7

Document S2. Tables S1-S14

## METHODS

### Generation of enhancer-reporter constructs

Genomic fragments spanning the *Runx1* enhancers of interest were generated by PCR (primer sequences in Table S1). Fragments were cloned downstream of a *LacZ or Gfp* gene in a reporter vector driven by the hsp68 core promoter^71^.

### Generation of transient transgenic embryos and transgenic mouse lines

Mouse transient transgenic embryos and transgenic mouse lines were generated by pronuclear injection of (CBA x C57BL/6)F1 zygotes following standard procedures. Transient transgenic embryos carrying *Runx1* enhancer-*LacZ* reporter construct were identified by *LacZ* specific PCR (primer sequences in Table S1) on genomic DNA isolated from yolk sac or ectoplacental cone biopsies. Transgenic founder mice carrying *Runx1* enhancer-*Gfp* reporter construct were identified by genomic PCR for *Gfp* on DNA isolated from tail or ear biopsies (primers in Table S1). Four transgenic 110GFP F0 founder mice were generated. Transgenic reporter gene expression was analysed in F1 embryos to identify those with GFP expression in a *Runx1*-specific pattern, and in line with the pattern observed in the corresponding enhancer-*LacZ* reporter transient transgenic embryos. A stable 110GFP transgenic line carrying <15 copies of the transgene was generated. All mice were housed with free access to food and water in an enhanced environment. All procedures were in compliance with United Kingdom Home Office regulations and the Oxford University Clinical Medicine Animal Welfare and Ethical Review Committee.

### Timed matings and embryo collection

For timed pregnancies, (CBA x C57BL/6)F1 females (5 –16 weeks old) were mated overnight with transgenic males (maintained on a mixed CBA x C57BL/6 background). Embryos were collected in CaCl_2_ and MgCl_2_-containing PBS (Gibco) supplemented with 10% FCS (Gibco), 50 U/ml penicillin, 50 mg/ml streptomycin (Cambrex Corporation). Embryos were staged by counting somite pairs (E8 to E10.5) or tail somite pairs (E11.5)^72^. 110GFP transgenic embryos were identified by fluorescence illumination (X-Cite 120; Improvision) on a Leica MZFLIII microscope.

### Analysis of enhancer-lacZ transient transgenic embryos

Transient transgenic embryos were analysed as described previously^30^. Briefly, embryos were fixed in PBS containing 1% formaldehyde, 0.2% glutaraldehyde, 2mM MgCl_2_, 5mM EGTA (pH 8.0), and 0.02% NP-40, washed in PBS 0.02% NP-40, and stained in Xgal staining solution (5mM K_3_Fe(CN)_6_, 5mM K_4_Fe(CN)_6_ 3 H_2_O, 2mM MgCl_2_, 0.01% sodium deoxycholate, 0.02% NP-40, and 1 mg/mL Xgal in PBS). Embryos were washed in PBS, postfixed, incubated in 15% sucrose in PBS, and embedded in Tissue-Tek OCT compound (Sakura, Siemens Medical Solutions Diagnostics). Transverse sections (8-10µm) were cut on a Leica CM3050S Cryostat (Leica Microsystems), collected on SuperFrost Plus microscope slides (VWR International) and coverslipped using Kaiser glycerol gelatin (VWR International). Embryo sections were examined, and pictures taken on a Nikon Eclipse E600 brightfield microscope with either a Nikon PlanFluor 40x/0.75 or 60x/1.40 objective and using the Nikon Digital Camera DXM 1200c setup.

### Dissections and generation of cell suspensions

Cell suspensions from dissected embryos were prepared as described^33^. Briefly, E8.5 concepti, E9.5 caudal parts (CP) and yolk sac (YS) and E10.5/E11.5 AGM with vitelline and umbilical arteries attached (AVU), were harvested and dissected in CaCl_2_ and MgCl_2_-containing PBS supplemented with 10% FCS (Gibco), 50U/ml penicillin, and 50µg/ml streptomycin (Cambrex Corporation) using forceps (Watchmakers) and 25G needles. Tissues were pooled and incubated in 0.12% (w/v) of collagenase (Type I, Sigma) in PBS without CaCl_2_ and MgCl_2_ supplemented with 10% FCS (Gibco), 50U/ml penicillin, and 50µg/ml streptomycin (Cambrex Corporation), dissociated by pipetting and washed. E14 fetal liver (FL) was dissociated by pipetting and washed. Viable cells were counted using Neubauer hemocytometer and trypan blue (Invitrogen) or NucleoCounter® NC-3000^TM^ automated cell counter (Chemometec).

### Flow cytometry and cell sorting

FACS analysis was performed as previously described^33^. In brief, embryonic cells were isolated using a 100µm nozzle on a BD FACSAria™ Fusion II instrument. Analysis was performed on CyAn ADP (Beckman Coulter) or BD LSRFortessa™ cell analyzer. Antibody staining was carried out in PBS (without CaCl_2_ and MgCl_2_) supplemented with 10% FCS (Gibco), 50U/ml penicillin, and 50µg/ml streptomycin (Cambrex Corporation). After sorting, cells were collected in tubes containing 100% FCS or sorted directly into OP9 co-culture media. Antibodies (Table S2) were titrated to determine their optimal concentration of use and gates were defined using unstained, single stained and fluorescence-minus-one stained cells. Dead cells were excluded based on Hoechst 33258 (Invitrogen) or 7AAD (eBioscience) uptake. Data were acquired and analysed using FACS Diva (BD) and FlowJo software.

### Immunofluorescence analysis and imaging

Sections for immunofluorescence analysis were processed as previously described^33^. In brief, embryos were fixed in 4% paraformaldehyde, washed in PBS and soaked in 15% (w/v) sucrose. Samples were snap-frozen in Tissue-Tek OCT compound (Sakura, Siemens Medical Solutions Diagnostics), sectioned at 10µm (Leica CM3050s) and collected onto glass slides (Superfrost plus, VWR). Sections were block permeated and stained in 5% FCS in 0.4% Triton-X/PBS (primary and secondary antibody list in Table S2). Slides were mounted with Vectashield hard-set mounting medium containing DAPI (Vector Laboratories). Image acquisition was performed using a Zeiss 510 and Zeiss 780 Inverted confocal microscopes. 25x (LD LCI PA 25x/0.8 DIC), 40x (LD C-Apochromat 40x/1.1) and 63x (Plan Apochromat 63x/1.4) objectives were used with oil immersion. Images were processed using Adobe Photoshop. Wholemount immunofluorescence was performed according to Yokomizo et al^73^. In brief, the head, tail, limbs and YS of E10.5 embryos were removed and embryos fixed in 4% paraformaldehyde and washed in PBS. For long-term storage, 50% and 100% MeOH washes were performed and embryos stored in 100% MeOH at −20°C. Prior to staining, embryos were washed in 50% MeOH and PBS and incubated in individual Eppendorf tubes in PBS 5-10% FCS 0.4% TritonX on a shaking rack. Primary and secondary staining (Table S2) were performed in 100-200μl of PBS 5-10% FCS 0.4% TritonX overnight at 4°C on a shaking rack. In between staining and after secondary antibody incubation, wholemounts were washed in the same buffer. Embryos were dehydrated in MeOH, cleared in BABB (1:2 benzyl alcohol:benzyl benzoate) and mounted into Fastwells (Grace Bio-Labs). Image acquisition was performed overnight using a Zeiss LSM 780 Upright confocal microscope with a 25x oil immersion objective (LD LCI PA 25x/0.8 DIC) and the ZEN 2011 LSM (black) software. Z-stacks and tiles of wholemount images were then 3D reconstructed in IMARIS (Bitplane) or OMERO software. Images were analysed with IMARIS (Bitplane) and Adobe Photoshop was used for final image processing.

### Hematopoietic progenitor assay

Colony-forming unit-culture (CFU-C) assays were performed using Methocult M3434 (Stem Cell Technologies). Cells were plated in duplicate in 35mm culture dishes according to manufacturer’s instructions. Cultures were grown at 37°C with 5% CO_2_ and colonies counted after 7 days.

### Long-term HSC reconstitution analysis

Male mice (C57BL/6xCBA)/F1 up to 3 months old were sub-lethally irradiated with a split dose of 9Gy from a 137Cs source. Donor cells were obtained from either E11.5 AVU or from OP9 co-aggregates and injected together with 2×10^5^ spleen carrier cells into the tail veins of irradiated recipients. The level of reconstitution in peripheral blood at 4 and 16 weeks post-transplant was determined by *Gfp*-specific PCR (Table S1) and quantitative analysis (Image Lab, Bio-Rad) of genomic DNA extracted from peripheral blood using the DNeasy® Blood & Tissue kit (Qiagen) according to manufacturer’s instructions. Percentage reconstitution was calculated based on comparison to standards containing 0%, 1%, 10%, 50%, and 100% of donor genomic DNA.

### OP9 maintenance and co-aggregate cultures

OP9 cells were maintained as previously described^28^. Sorted cell populations from pools of E10.5/E11.5 AVU were assayed as previously described^12,74,75^. In brief, a cell mixture containing AGM cells (1ee) and 10^5^ OP9 was suspended in Iscove’s modified Dulbecco’s medium (IMDM, Gibco), aspirated in a 200ml pipette tip sealed with paraffin, and centrifuged at 400*g* for 5 min at RT. Co-aggregates were transferred onto a floating 0.8μm nitrocellulose membranes (Millipore) and cultured at the liquid–gas interface in IMDM (Gibco) supplemented with 100ng/ml IL-3, 100ng/ml SCF, 100ng/ml Flt3 ligand (Peprotech), L-glutamine and penicillin/streptomycin (Gibco). After 5 days, aggregates were dissociated and analysed as previously described^12^.

### Single-cell RNA-seq cell isolation and library preparation

AVU from pooled E10 23GFP Tg (9 embryos, 32-35 sp) and 110GFP Tg (5 embryos, 32-34 sp) were dissected and processed into single cell suspension as described above. Cells were incubated in TruStain FcX™ PLUS blocking reagent (Biolegend) and TotalSeqB antibody cocktail (Biolegend) (Table S2), as per manufacturer’s instructions. 23GFP+ and 110GFP+ cells were FACS-sorted using a BD FACSAria Fusion sorter (100μm nozzle) and cell viability was assessed (99% and 95% live cells for 23GFP and 110GFP, respectively) before loading each sample onto independent channels of a Chromium chip for single cell partitioning and barcoding on Chromium Controller (10x Genomics). Single-cell cDNA synthesis and library preparation were performed using a Chromium Next GEM Single Cell 3’ v3.1 Dual Index Kit (10x Genomics) as per manufacturer’s instructions and sequenced on Illumina NovaSeq 6000.

### Single-cell RNA-seq analysis

Raw sequencing data were processed using Cell Ranger Software (version 5.0.0, 10x Genomics) and a customised version of the GRCm39 mouse genome assembly where the sequence for the *Gfp* gene was added using the cellranger mkref pipeline (10x Genomics). After sequencing, 5,751 cells were recovered for 23GFP AVU and 2,997 for 110GFP AVU. Ambient RNA, mostly consisting in low expression of globin genes, was removed using SoupX package^76^ (contamination fraction was selected manually as 10% for 23GFP AVU and 20% for 110GFP AVU). Low quality cells and empty droplets were filtered out based on the number of detected genes, UMI and percentage of mitochondrial genes (< 5%) and doublets were identified and excluded using Scrublet^77^. Downstream analysis was performed using Seurat package (version 4.0.4)^78^ in R (version 4.1.3). Each sample was normalised and scaled independently using SCTransform^79^ (including cell cycle-associated genes, UMI counts and mitochondrial genes regression) and samples were integrated using Seurat RPCA approach (k=5). When Seurat subset function was used to remove contaminating clusters or select specific populations in the original dataset, SCTransform and RPCA integration were performed again on the selected subsets using the same parameters. To improve cluster annotation, MAGIC^80^ was used to impute missing values and reduce dropout effects. Clusters were annotated based on differentially expressed gene (DEG) analysis (using Seurat FindAllMarkers, min.pct = 0.25, logfc.threshold = 0.25) and the expression of known cell type-specific marker genes. The FindSubCluster function was used when marker gene expression suggested further heterogeneity within a defined cluster. Finally, cluster annotation was confirmed by visualising the expression of gene signatures from publicly available datasets including embryonic hematopoietic tissues^37–39,42,81^ using the AddModuleScore function (ctrl = 100).

### HSC potential gene signature

To identify marker genes specifically expressed in HSC^Pot^ when compared to HE and HSC^Not^, DEG analysis was performed using Seurat FindAllMarkers function (3-way comparison) and only genes that showed a minimum difference of 25% in the fraction of detection (min.diff.pct = 0.25) and an average log2 fold-change of at least 0.5 (logfc.threshold = 0.5) between groups were tested.

### Single-cell Multiome cell isolation and library preparation

AVU from pooled E9 (33 embryos, 20-25 sp) and E10 (24 embryos, 32-36 sp) 23GFP Tg were dissected, processed into single cell suspension and stained for FACS sorting, as described above. Ter119-23GFP+ cells were FACS-sorted using a BD FACSAria Fusion sorter (100μm nozzle) and nuclei were isolated as 10x Genomics demonstrated protocols (CG000365). Nuclei quality was assessed, and nuclei were counted before loading each sample onto independent channels of a Chromium chip for single cell partitioning and barcoding on Chromium Controller (10x Genomics). Single-cell cDNA synthesis and library preparation were performed using a Chromium Next GEM Single Cell Multiome ATAC + Gene Expression Kit (10x Genomics) as per manufacturer’s instructions and sequenced on Illumina NovaSeq 6000.

### Single-cell Multiome analysis

Raw sequencing data were processed using Cell Ranger ARC (version 2.0.2, 10x Genomics) and GRCm38 mouse genome assembly. Downstream analysis was performed using Signac (version 1.12.0)^82^ and Seurat packages (version 5.0.1)^78^ in R (version 4.3.2). Per-cell quality control metrics were computed on the merged samples for both RNA (1000 < detected genes < 7000, 1000 < UMI< 70000 and mitochondrial genes < 20%) and ATAC (2.5 < TSS enrichment score < 10, Nucleosome signal < 1.5 and 1500 < ATAC fragments < 25000) assays, and low-quality nuclei were filtered out. Gene expression data of the 7349 nuclei that passed QC were normalised and scaled using SCTransform^79^ (including cell cycle-associated genes, UMI counts and mitochondrial genes regression). Clusters were identified using the FindClusters function (resolution = 1) in Seurat and dimension reduction was performed using RunUMAP function (dims = 1:50). Clusters were annotated based on differentially expressed gene (DEG) analysis (using Seurat FindAllMarkers, min.pct = 0.25, logfc.threshold = 0.25) and the expression of known cell type-specific marker genes. ATAC-seq peaks were identified using MACS2^83^ and the CallPeaks function in Signac (additional.args = “-q 0.01”). DNA accessibility data were processed by performing latent semantic indexing (LSI) in Signac and dimension reduction was performed using the RunUMAP function (reduction = LSI, dims= 2:50). Joint UMAP visualisation of RNA and ATAC assays was computed using the weighted nearest neighbour method (WNN) in Seurat. When Seurat subset function was used to select *Hoxa+* (*hoxa7*, *hoxa9* and *hoxa10*) hematopoietic clusters in the original dataset, both RNA and ATAC processing was performed again using the same parameters. Peak-to-gene linkage on analysis on *Hoxa+* clusters was performed using the LinkPeaks function in Signac. HSC-fated signature was calculated as described above.

### Motif analysis

Genomic regions corresponding to the ATAC peaks of the *Runx1* +3 and +110 enhancers were searched for known transcription factor motifs using FIMO^84^ from the MEME suite (version 5.5.3) and the HOCOMOCOv11_core_MOUSE database^85^.

Mean conservation score for each motif was calculated by intersecting their sequence with base wise conservation scores (phyloP) of 59 vertebrate genomes with mouse GRCm38 from UCSC, using Bedtools (version 2.29.2). Significant hits resulting from FIMO analysis were then filtered by *i)* mean conservation score > 0; *ii)* presence within the highly conserved region of each enhancer (i.e. conserved in mouse, human, dog and opossum) and *iii)* expression in the cell type of interest.

### Statistical analysis

Results are presented as average ± SD. Unpaired, two-tailed Student’s t-test with Welch’s correction was used to determine the level of significance. The following p-values were considered statistically significant and were highlighted by asterisks: * p<0.05; ** p<0.005.

