## Supplementary Information for "Resolving hematopoietic stem versus progenitor cell potential in the mouse dorsal aorta by differential *Runx1* +110 enhancer activity"

#### SUPPLEMENTAL INFORMATION

### Figure S1

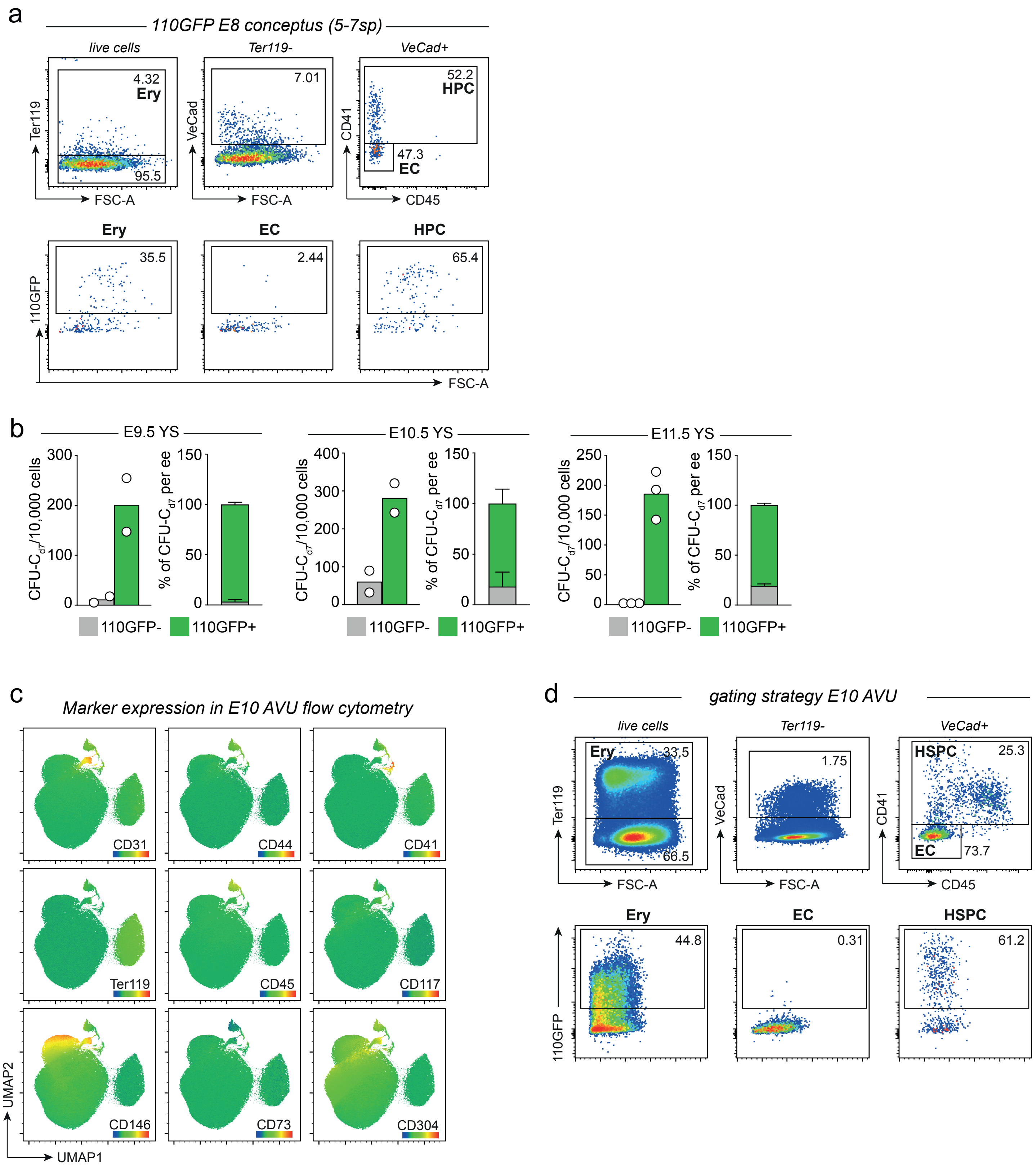

a
110GFP E8 conceptus (5-7sp)
live cells
Ter119-
VeCad+
Ter119
VeCad
CD41
FSC-A
CD45
4.32
Ery
95.5
7.01
52.2
HPC
47.3
EC
Ery
EC
HPC
110GFP
35.5
2.44
65.4
b
E9.5 YS
E10.5 YS
E11.5 YS
CFU-C d7/10,000 cells
% of CFU-C d7 per ee
110GFP-
110GFP+
c
Marker expression in E10 AVU flow cytometry
CD31
CD44
CD41
Ter119
CD45
CD117
CD146
CD73
CD304
UMAP2
UMAP1
d
gating strategy E10 AVU
live cells
Ter119-
VeCad+
Ery
33.5
66.5
1.75
HSPC
25.3
EC
73.7
Ery
EC
HSPC
44.8
0.31
61.2
110GFP

**Figure S1. The *Runx1* +110 mediates reporter gene expression to distinct subsets of *Runx1*-expressing hematopoietic cells at the onset of hematopoiesis (related to Figure 1).**

**a.** Gating strategy of flow cytometry analysis of E8 (5-7sp) whole conceptus of 110GFP transgenic embryos and representative plots showing GFP+ expression in each cell type.

**b.** CFU-C potential of 110GFP+ and 110GFP- cells isolated from YS of E9.5 (19-25sp), E10.5 (32-35sp) and E11.5 (14-17tsp) transgenic embryos. Absolute number of CFU-$C_{d7}$ per 10,000 cells and frequency of CFU-$C_{d7}$ per embryo equivalent (ee) are reported (Mean ± SD of n=2 for E9.5, n=2 for E10.5 and n=3 for E11.5 independent experiments).

**c.** Mean fluorescence intensity (MFI) of flow cytometry markers used for cell type annotation of E10.5 AVU sample in Fig. 1f.

**d.** Gating strategy of flow cytometry analysis of E10.5 AVU and representative plot showing GFP+ expression in 110GFP transgenic embryos.

### Figure S2

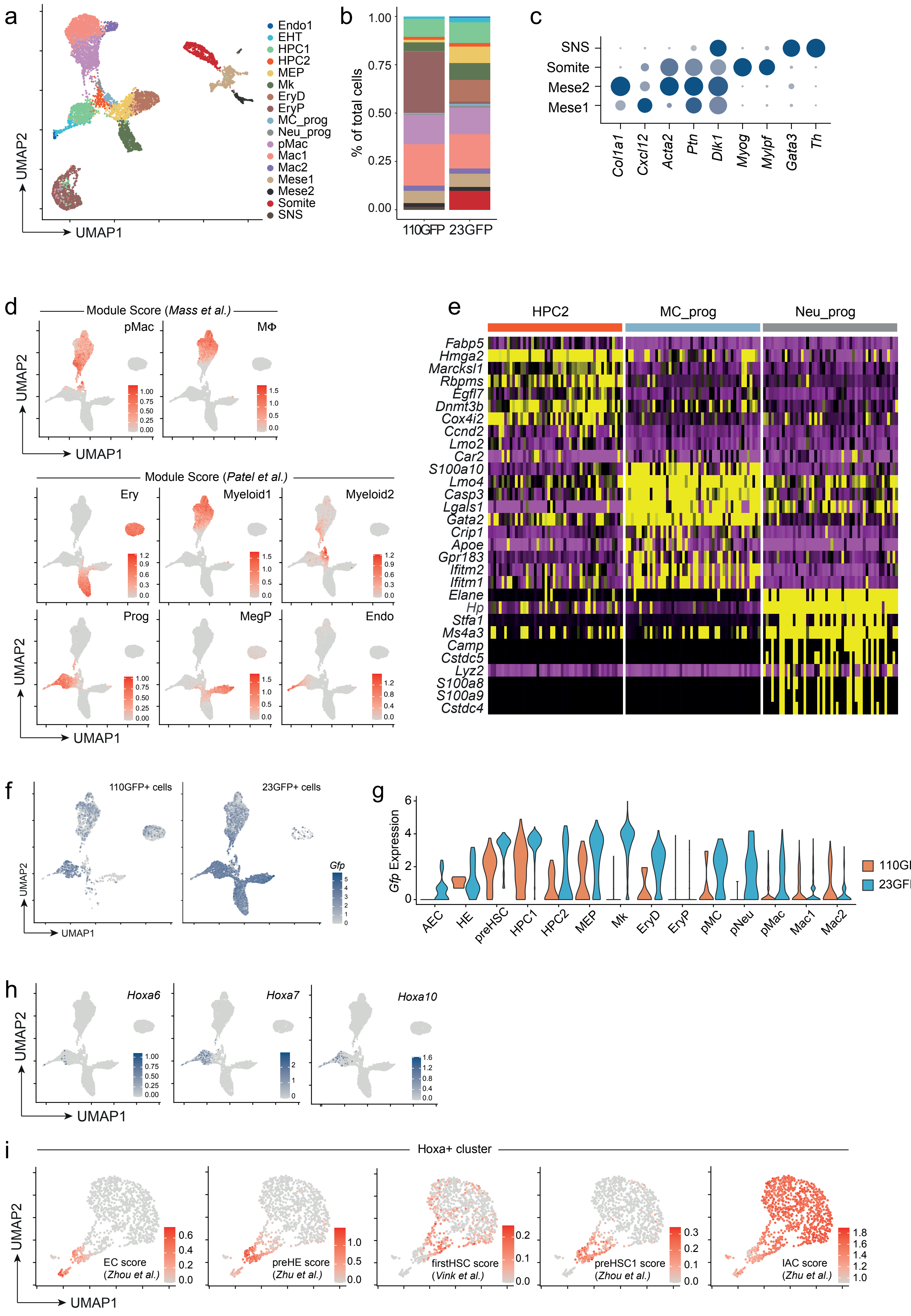
a
Endo1
EHT
HPC1
HPC2
MEP
Mk
EryD
EryP
MC_prog
Neu_prog
pMac
Mac1
Mac2
Mese1
Mese2
Somite
SNS
UMAP1
UMAP2
b
% of total cells
110GFP
23GFP
c
SNS
Somite
Mese2
Mese1
Col1a1
Cxcl12
Acta2
Ptn
Dlk1
Myog
Mylpf
Gata3
Th
d
Module Score (Mass et al.)
pMac
MΦ
Module Score (Patel et al.)
Ery
Myeloid1
Myeloid2
Prog
MegP
Endo
e
HPC2
MC_prog
Neu_prog
Fabp5
Hmga2
Marcksl1
Rbpms
Egfl7
Dnmt3b
Cox4i2
Ccnd2
Lmo2
Car2
S100a10
Lmo4
Casp3
Lgals1
Gata2
Crip1
Apoe
Gpr183
Ifitm2
Ifitm1
Elane
Hp
Stfa1
Ms4a3
Camp
Cstdc5
Lyz2
S100a8
S100a9
Cstdc4
f
110GFP+ cells
23GFP+ cells
Gfp
g
Gfp Expression
AEC
HE
preHSC
HPC1
HPC2
MEP
Mk
EryD
EryP
pMC
pNeu
pMac
Mac1
Mac2
110GFP
23GFP
h
Hoxa6
Hoxa7
Hoxa10
i
Hoxa+ cluster
EC score
(Zhou et al.)
preHE score
(Zhu et al.)
firstHSC score
(Vink et al.)
preHSC1 score
(Zhou et al.)
IAC score
(Zhu et al.)

**Figure S2. The 110GFP transgene marks primitive macrophages, EryP and emerging *Hoxa*+ HSPCs (related to Figure 2).**

- **a.** UMAP representation of scRNAseq profiling of E10.5 AVU of 110GFP (2,997 cells) and 23GFP (5,751 cells) transgenic embryos before excluding non-hematopoietic contaminants. Mese1 and 2, Mesenchyme1 and 2; SNS, sympathetic nervous system.
- **b.** Bar graph quantification displaying frequency of clusters within each sample.
- **c.** Dot plot of marker genes used to annotate non-hematopoietic clusters in scRNAseq data in Fig. S4a.
- **d.** Expression of gene signatures from publicly available datasets including macrophage precursors (pMac), embryonic macrophages[61] and E11 AGM- and FL-derived hematopoietic cells[41].
- **e.** Heatmap of DEG comparing Hoxa- progenitors isolated from E10 AVU of 110GFP and 23GFP transgenic embryos. HPC, hematopoietic progenitor cell 2; MC_prog, mast cell progenitor; Neu_prog, neutrophil progenitor.
- **f.** Expression of *Gfp* reporter gene in 110GFP+ (left) and 23GFP+ (right) integrated UMAP.
- **g.** Violin plot displaying levels of expression of *Gfp* reporter gene in each UMAP cluster, split by sample of origin (110GFP orange and 23GFP light blue).
- **h.** Expression of *Hoxa6, Hoxa7* and *Hoxa10* in 110GFP+ and 23GFP+ integrated dataset (Figure 2b).
- **i.** Expression of gene signatures from publicly available datasets including embryonic endothelial cells (EC)[36], first HSC[38], preHE and IAC[37].

### Figure S3

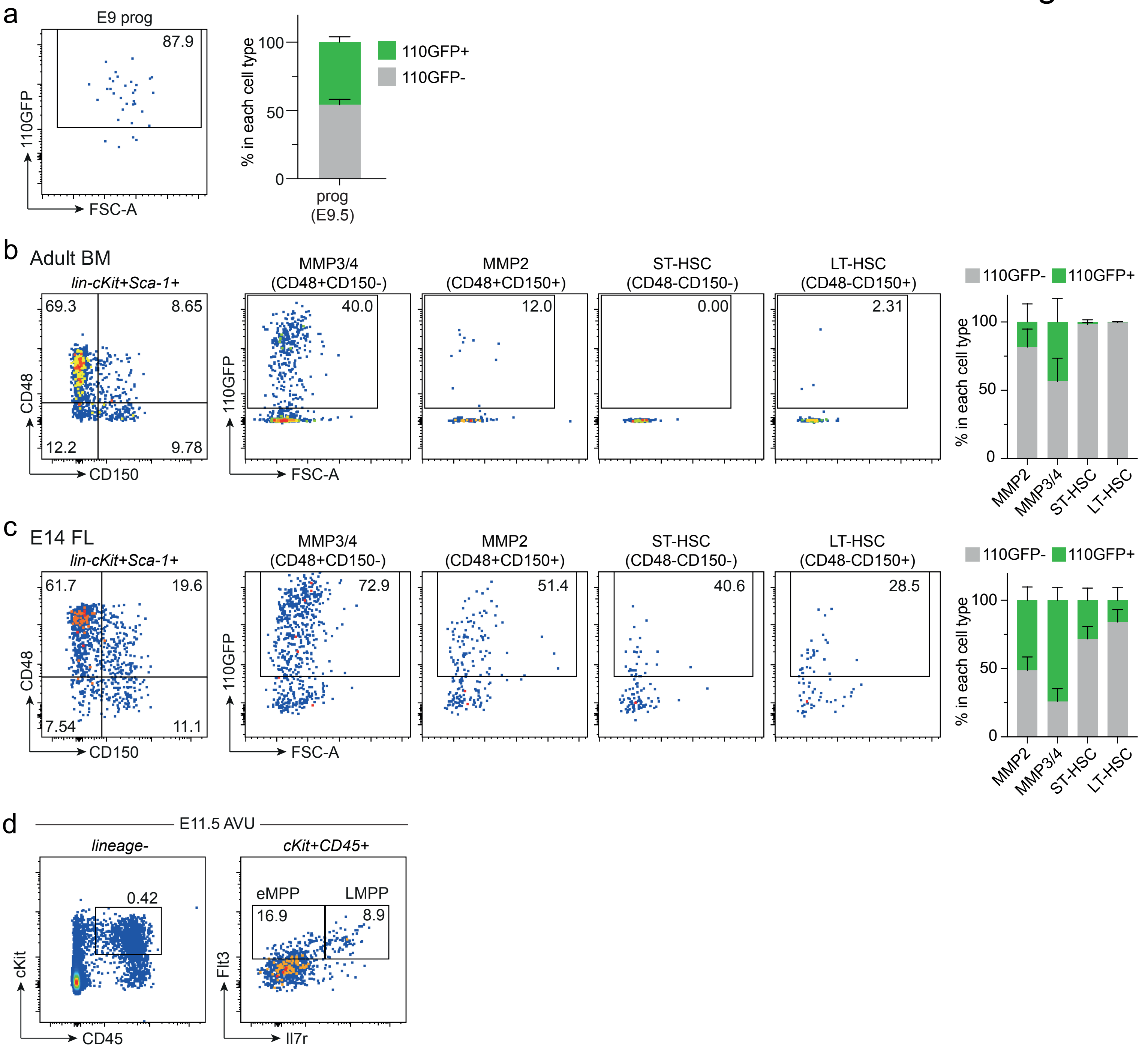

a
E9 prog
87.9
110GFP
FSC-A
% in each cell type
100
50
0
110GFP+
110GFP-
prog
(E9.5)
b
Adult BM
lin-cKit+Sca-1+
69.3
8.65
12.2
9.78
CD48
CD150
MMP3/4
(CD48+CD150-)
40.0
MMP2
(CD48+CD150+)
12.0
ST-HSC
(CD48-CD150-)
0.00
LT-HSC
(CD48-CD150+)
2.31
110GFP
FSC-A
110GFP-
110GFP+
% in each cell type
100
50
0
MMP2
MMP3/4
ST-HSC
LT-HSC
c
E14 FL
lin-cKit+Sca-1+
61.7
19.6
7.54
11.1
CD48
CD150
MMP3/4
(CD48+CD150-)
72.9
MMP2
(CD48+CD150+)
51.4
ST-HSC
(CD48-CD150-)
40.6
LT-HSC
(CD48-CD150+)
28.5
110GFP
FSC-A
110GFP-
110GFP+
% in each cell type
100
50
0
MMP2
MMP3/4
ST-HSC
LT-HSC
d
E11.5 AVU
lineage-
0.42
cKit
CD45
cKit+CD45+
eMPP
LMPP
16.9
8.9
Flt3
Il7r

**Figure S3. Cells belonging to the functional HSC lineage are not marked by the 110GFP reporter (related to Figure 3).**

- **a.** Representative flow cytometry plot of 110GFP expression in E9.5 progenitors (prog). Mean ± SD of pooled PAS from n=3 independent experiments.
- **b.** Gating strategy to assess 110GFP expression in phenotypically defined ST-HSCS (CD48-CD150- LSK), LT-HSCS (CD48-CD150+ LSK), MMP2 (CD48+CD150+ LSK) and MMP3/4 (CD48+CD150- LSK). Frequency ± SD of n=3 adult bone marrow.
- **c.** Gating strategy to assess 110GFP expression in phenotypically defined ST-HSCS (CD48-CD150- LSK), LT-HSCS (CD48-CD150+ LSK), MMP2 (CD48+CD150+ LSK) and MMP3/4 (CD48+CD150- LSK). Frequency ± SD of n=23 E14 FL.
- **d.** Gating strategy to assess 110GFP expression in phenotypically defined eMPP (lin-CD45+cKit+Flt3+Il7r-) and LMPP (lin-CD45+cKit+Flt3+Il7r+) from E11.5 (15-20tsp) AVU.

Figure S4

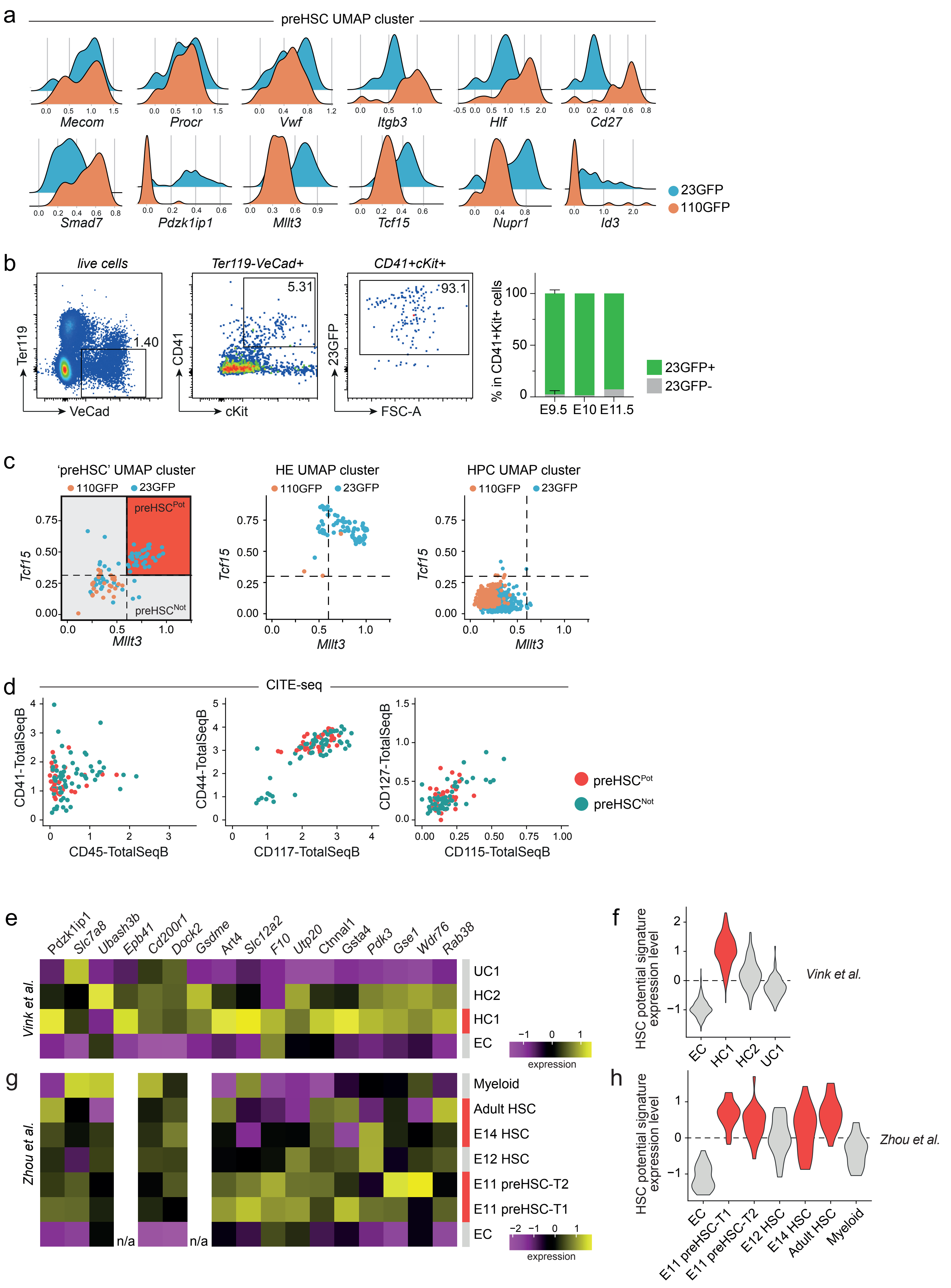

a
preHSC UMAP cluster
Mecom
Procr
Vwf
Itgb3
Hlf
Cd27
Smad7
Pdzk1ip1
Mllt3
Tcf15
Nupr1
Id3
23GFP
110GFP
b
live cells
Ter119-VeCad+
CD41+cKit+
Ter119
VeCad
1.40
CD41
cKit
5.31
23GFP
FSC-A
93.1
% in CD41+Kit+ cells
100
50
0
E9.5
E10
E11.5
23GFP+
23GFP-
c
'preHSC' UMAP cluster
HE UMAP cluster
HPC UMAP cluster
110GFP
23GFP
preHSC^Pot
preHSC^Not
Tcf15
Mllt3
d
CITE-seq
CD41-TotalSeqB
CD45-TotalSeqB
CD44-TotalSeqB
CD117-TotalSeqB
CD127-TotalSeqB
CD115-TotalSeqB
preHSC^Pot
preHSC^Not
e
Pdzk1ip1
Slc7a8
Ubash3b
Epb41
Cd200r1
Dock2
Gsdme
Art4
Slc12a2
F10
Utp20
Ctnnal1
Gsta4
Pdk3
Gse1
Wdr76
Rab38
Vink et al.
UC1
HC2
HC1
EC
expression
f
HSC potential signature expression level
Vink et al.
EC
HC1
HC2
UC1
g
Zhou et al.
Myeloid
Adult HSC
E14 HSC
E12 HSC
E11 preHSC-T2
E11 preHSC-T1
EC
n/a
n/a
expression
h
HSC potential signature expression level
Zhou et al.
EC
E11 preHSC-T1
E11 preHSC-T2
E12 HSC
E14 HSC
Adult HSC
Myeloid

**Figure S4. Emergence of HSC potential in 110GFP- cells is marked by a 17-gene signature and expression of genes associated with epigenetic repression (related to Figure 4).**

- **a.** Histograms comparing expression of known marker genes associated with embryonic HSPCs and adult LT-HSCs in transcriptionally defined preHSCs UMAP cluster. Cells coloured based on original sample (110GFP orange and 23GFP light blue).
- **b.** Gating strategy and quantification of flow cytometry data measuring the expression of 23GFP transgene in Ter119-VeCad+CD41+cKit+ cells in n=4 E9.5 (19-28sp) PAS, n=1 E10 (29-31sp) and n=1 E11.5 AVU.
- **c.** Scatter plots illustrating the approach used to refine the annotation of single cells in the preHSC UMAP cluster based on (pre)HSC potential. Cells are coloured based on sample of origin (110GFP orange and 23GFP light blue). Red quadrant highlights functionally defined preHSC$^{Pot}$. Expression of *Tcf15* and *Mllt3* was compared to transcriptionally defined HE and HPC UMAP clusters.
- **d.** Scatter plots of CITE-seq data displaying the expression of common hematopoietic surface markers in preHSC cluster. Cells coloured based on functional analysis (preHSC$^{Not}$ teal and preHSC$^{Pot}$ red).
- **e.** Heatmap displaying the expression of each single gene included in the HSC potential 17-gene signature in functionally and transcriptionally defined HSC from Vink et al.[37]. Each row represents a pseudobulk average of all cells from a single cluster. Functionally defined HSC clusters are coloured in red.
- **f.** Expression of the HSC potential signature (17 genes) in functionally and transcriptionally defined HSC from Vink et al.[37]. Functionally defined HSC clusters are coloured in red.
- **g.** Heatmap displaying the expression of each single gene included in the HSC potential 17-gene signature in functionally and transcriptionally defined HSCs from Zhou et al.[35]. Each row represents a pseudobulk average of all cells from a single cluster. Functionally defined HSC clusters are coloured in red.
- **h.** Expression of the HSC potential signature (17 genes) in functionally and transcriptionally defined HSCs from Zhou et al.[35]. Functionally defined HSC clusters are coloured in red.

### Figure S5

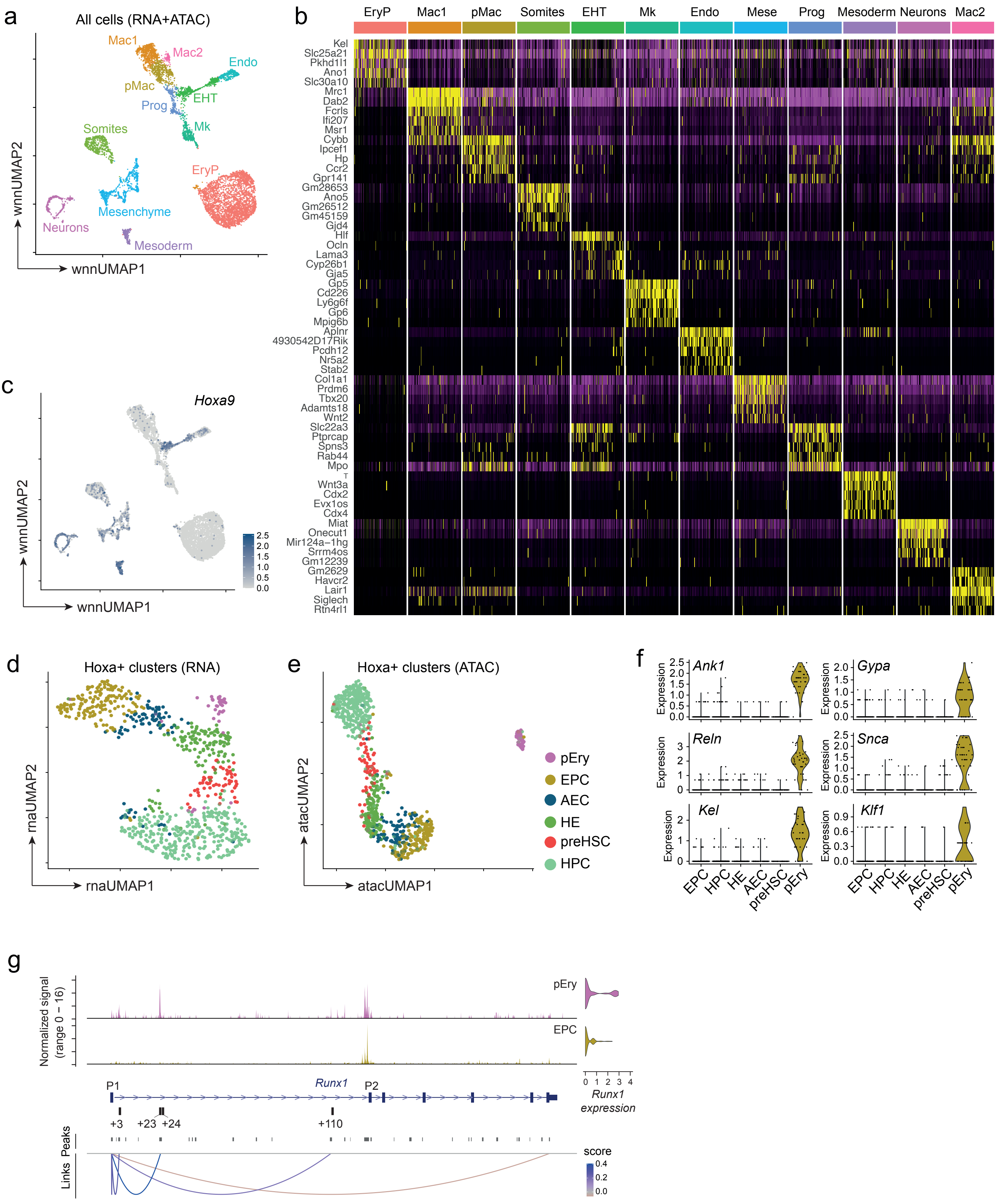

a
All cells (RNA+ATAC)
Mac1
Mac2
Endo
pMac
EHT
Prog
Mk
Somites
EryP
Mesenchyme
Neurons
Mesoderm
wnnUMAP2
wnnUMAP1
b
EryP Mac1 pMac Somites EHT Mk Endo Mese Prog Mesoderm Neurons Mac2
Kel
Slc25a21
Pkhd1l1
Ano1
Slc30a10
Mrc1
Dab2
Fcrls
Ifi207
Msr1
Cybb
Ipcef1
Hp
Ccr2
Gpr141
Gm28653
Ano5
Gm26512
Gm45159
Gjd4
Hlf
Ocln
Lama3
Cyp26b1
Gja5
Gp5
Cd226
Ly6g6f
Gp6
Mpig6b
Aplnr
4930542D17Rik
Pcdh12
Nr5a2
Stab2
Col1a1
Prdm6
Tbx20
Adamts18
Wnt2
Slc22a3
Ptprcap
Spns3
Rab44
Mpo
T
Wnt3a
Cdx2
Evx1os
Cdx4
Miat
Onecut1
Mir124a−1hg
Srrm4os
Gm12239
Gm2629
Havcr2
Lair1
Siglech
Rtn4rl1
c
Hoxa9
2.5
2.0
1.5
1.0
0.5
0.0
wnnUMAP2
wnnUMAP1
d
Hoxa+ clusters (RNA)
rnaUMAP2
rnaUMAP1
e
Hoxa+ clusters (ATAC)
atacUMAP2
atacUMAP1
pEry
EPC
AEC
HE
preHSC
HPC
f
Ank1
Gypa
Reln
Snca
Kel
Klf1
Expression
EPC HPC HE AEC preHSC pEry
g
pEry
EPC
Normalized signal (range 0 – 16)
0 1 2 3 4
Runx1 expression
P1
P2
Runx1
+3
+23 +24
+110
Peaks
Links
score
0.4
0.2
0.0

**Figure S5. A single-cell multiomic datset of gene expression and DNA accessibility in Ter119-23GFP+ cells from E9 and E10 AVUs (related to Figure 4).**

- **a.** UMAP representation of joint RNA and ATAC measurements using weighted nearest neighbour (WNN) integration for all clusters from multiomic data.
- **b.** Heatmap of DEG comparing all clusters in RNA assay of scMultiome data. EryP, primitive erythrocytes; Mac1, macrophage 1; pMac, macrophage precursors; EHT, endothelial-to-hematopoietic transition; Mk, megakaryocytes; Endo, endothelial cells; Mese, mesenchymal cells; Prog, hematopoietic progenitors; Mac2, macrophage 2.
- **c.** Expression of *Hoxa9* in RNA assay of scMultiome data (all clusters).
- **d.** UMAP representation of RNA measurements in *Hoxa*+ clusters from multiomic data.
- **e.** UMAP representation of ATAC measurements in *Hoxa*+ clusters from multiomic data.
- **f.** Expression of erythrocyte-associated genes in *Hoxa*+ clusters from multiomic data.
- **g.** ATAC-seq tracks and peak-gene links for the *Runx1* locus in *Hoxa*+ clusters. pEry, erythroid progenitor; EPC, endothelial progenitor cell.

Figure S6

#### Runx1 +3 enhancer

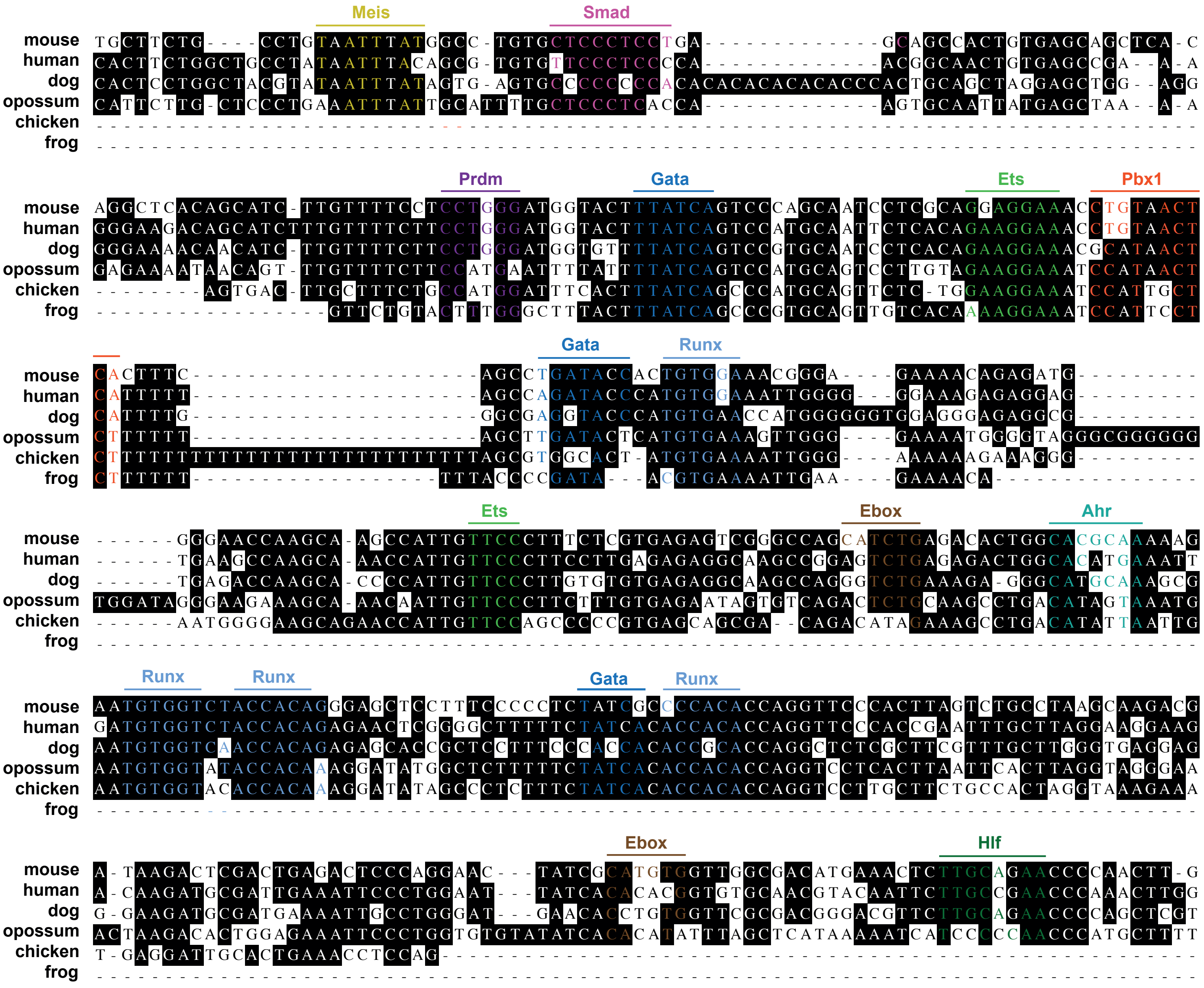

**Figure S6. Multispecies alignment of the highly conserved phylogenetic region of the Runx1 +3 enhancer (related to Figure 4).**

Nucleotides highlighted in black are conserved between all species analysed. Matches to consensus transcription factor binding sites (TFBS) in phylogenetic footprints are highlighted (Meis, yellow; Smad, pink; Prdm, purple; Gata, dark blue; Ets, light green; Pbx1, red; Runx, light blue; Ahr, teal; Hlf, dark green; Ebox, brown).

Figure S7

#### Runx1 +110 enhancer

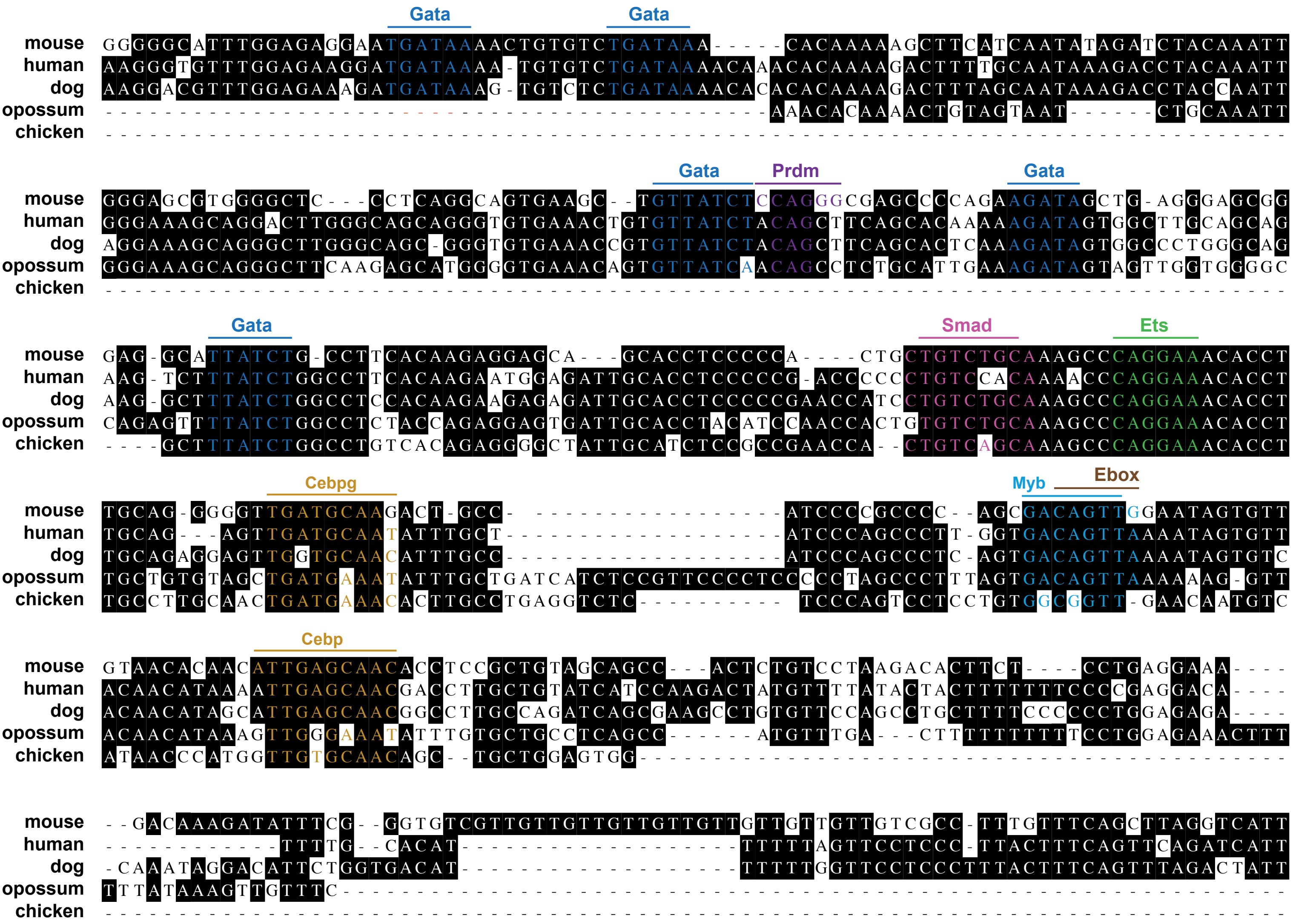

**Figure S7. Multispecies alignment of the highly conserved phylogenetic region of the Runx1 +110 enhancer (related to Figure 4).**

Nucleotides highlighted in black are conserved between all species analysed. Matches to consensus transcription factor binding sites (TFBS) in phylogenetic footprints are highlighted (Gata, dark blue; Prdm, purple; Smad, pink; Ets, light green; Cebp, gold; Myb, light blue; Ebox, brown).
